# A fast-to-faithful transition shapes the DNA repair landscape during embryogenesis

**DOI:** 10.64898/2026.04.22.720229

**Authors:** Shengzhou Wang, Ian Pantziris, Mengsheng Zhang, Yang Liu, James A. Gagnon

## Abstract

Accurate and timely repair of DNA double-strand breaks (DSBs) is essential for genome maintenance in all cells. Embryos are particularly vulnerable to DSBs. During zebrafish development, a single fertilized cell undergoes rapid divisions to form an embryo of 50,000 cells in the first 24 hours, subjecting its genome to intense replication stress and the inevitable formation of genomic DSBs. While we know that failure to repair these breaks can result in embryonic lethality, the kinetics, fidelity, and pathway choice of DSB repair during embryogenesis are not well understood. Here, we used light-activated CRISPR to generate targeted genomic DSBs across zebrafish embryo development. Importantly, DSB induction occurs within seconds after light stimulation, enabling precise measurements of repair kinetics within a single cell cycle. We found that DSBs were repaired within 15 minutes during the early, rapid-division stages. At later stages, the pace of DNA repair declines as the cell cycle slows. By leveraging mathematical modeling and mutants that disrupt DNA repair pathways, we uncovered a developmental transition from error-prone microhomology-mediated end joining to more faithful non-homologous end joining that correlates with the gradual shift in repair kinetics. To our knowledge, this is the first study to resolve DSB repair dynamics with high temporal resolution during embryo development. Our study establishes a framework for systematically interrogating the cellular responses to DNA damage in living model organisms.

**SIGNIFICANCE STATEMENT:** This study reveals how developing embryos rapidly repair dangerous DNA breaks while balancing speed and accuracy. Using a light-activated CRISPR system, the authors precisely control when DNA damage occurs and measure repair in real time in living zebrafish embryos. They show that early embryos prioritize extremely fast but error-prone repair, then transition to slower, more accurate mechanisms as development progresses. This “fast-to-faithful” shift explains how embryos protect their genomes during rapid cell division. The work provides a new framework for studying DNA repair in vivo and has implications for understanding developmental disorders, genome editing outcomes, and how cells manage DNA damage under changing biological conditions.

## INTRODUCTION

DNA double-strand breaks (DSBs) are the most deleterious form of genomic damage. DSBs can result from endogenous replication/transcription stress or exogenous insults like radiation, pollutants, and chemotherapeutics (Shrivastav et al. 2008; Ceccaldi et al. 2016). If incorrectly repaired, DSBs can cause mutations and chromosomal rearrangements. Unrepaired or sustained DSBs promote catastrophic chromosome loss (Zuccaro et al. 2020; Nahmad et al. 2022) and cell death (Krenning et al. 2019). To prevent genome instability and associated consequences, eukaryotic cells have evolved several pathways for repairing DSBs, including homologous recombination (HR), classical nonhomologous DNA end-joining (NHEJ), and microhomology-mediated end-joining (MMEJ) pathways (Ceccaldi et al. 2016; Kumari et al. 2025).

While DNA repair mechanisms are extensively characterized in cultured cells and animals (Jackson and Bartek 2009; Scully et al. 2019), far less is known about how these pathways operate in developing embryos. Developing embryos of many species face particular challenges for DNA repair. Their cells are often rapidly dividing, providing less time to identify and resolve DSBs. The stakes for the embryo are high - mistakes in DNA repair in an embryonic cell will have grave consequences for all descendent cells. The zebrafish embryo presents an illustrative example. Upon fertilization, the embryo employs rapid, synchronous rounds of DNA replication and cell division every 15 minutes, bypassing G1 and G2 phases for the first three hours (Kimmel et al. 1995; Siefert et al. 2015). The zygotic genome is transcriptionally silent and DNA damage checkpoints are inactive during this period (Vastenhouw et al. 2019). As development progresses, the rate of cell division slows and becomes asynchronous. Cells begin gene expression and acquire full cell cycle regulation, including the ability to arrest in response to DNA damage (Kermi et al. 2019; Brantley and Di Talia 2021). After 24 hours, the zebrafish embryo is composed of ∼50,000 cells, spanning dozens of cell types (Wagner et al. 2018; Sur et al. 2023). How can embryonic cells in this rapidly growing organism repair DSBs rapidly and accurately to avoid passing mutations to descendants that cause developmental disorders or embryonic lethality? How do DNA repair pathways adapt to the dynamic cellular landscape of the embryo to maintain genome integrity?

Technical limitations have hindered the study of DSB repair dynamics in living embryos. Early studies using high-energy irradiation to create DNA damage suggested that NHEJ is essential for cell survival in zebrafish embryos 6 hours post-fertilization (Bladen et al. 2005, 2007). In these cases, DSBs were induced randomly across the genome without sequence specificity, making it impossible to determine repair outcomes and contribution from other repair pathways. Subsequent work using exogenous fluorescence reporters and meganucleases as DSB inducers allowed for studying DSB repair at a specific site, discovering that all three major pathways are active during early development, with MMEJ and NHEJ showing particularly much stronger activity than HR (Liu et al. 2012; He et al. 2015). More recently, CRISPR-based approaches revealed that MMEJ dominates genomic DSB repair in the early embryo (Thyme and Schier 2016; Carrara et al. 2025). However, these approaches all generate asynchronous DSBs and are restricted to studies in the earliest stages of embryogenesis. Because of these limitations, it remains unclear how DSB repair can accommodate the fast cell divisions of early embryo development, whether repair kinetics are coordinated with changes in embryo cell cycle, or how the relative contributions of each pathway influence repair outcomes throughout embryogenesis.

Here we adapted a light-activated very fast (vf)CRISPR system (Liu et al. 2020) to enable time-resolved investigation of DSB repair dynamics across embryogenesis. Using this tool, we induced DSBs in zebrafish embryos at different developmental stages and monitored DNA repair kinetics and outcomes. We found that DSBs are repaired within minutes of their induction in the early embryo. DSB repair rate slows dramatically at later stages, in parallel with the slowing of the cell cycle, and repair outcomes change. Using mathematical modeling and mutants that disrupt MMEJ or NHEJ, we found that these shifts in repair kinetics and outcomes are dependent on a developmental transition from MMEJ- to NHEJ-mediated repair. Together, our study delineates the rapid pace and shifting dynamics of DNA repair in the developing zebrafish embryo.

## RESULTS

### vfCRISPR induces DSBs in zebrafish embryos within seconds

Studying DSB repair during embryogenesis requires tools that create genomic DSBs precisely, rapidly, and at different stages of development. Previous work demonstrated light activation of CRISPR-Cas9 targeting specific genes in zebrafish embryos using caged guide RNAs (cgRNAs) (Moroz-Omori et al. 2020; Zhou et al. 2020). Although efficient, these approaches lack temporal precision for DSB induction and are better suited to gene editing applications. In these methods, *Streptococcus pyogenes* Cas9 complexes with cgRNA to form ribonucleoproteins (RNPs) that remain inactive and diffuse throughout the nucleus. Only after light-mediated uncaging, can RNPs begin searching for and binding their genomic target sites to initiate DSBs - a process that takes tens of minutes in bacterial and human cells (Jones et al. 2017; Rose et al. 2017), and longer than the cell cycle of early zebrafish embryos.

To enable precise control of DSB induction, we adapted vfCRISPR, a Cas9 system that initiates targeted DSBs seconds after light stimulation (Liu et al. 2020). Combining vfCRISPR with time-resolved imaging and sequencing techniques has revealed a variety of fast processes during DSB repair in cultured cells (Liu et al. 2020; Zou et al. 2022; He et al. 2024; Marin-Gonzalez et al. 2025). vfCRISPR uses wild-type Cas9 protein and cgRNAs chemically modified with light-sensitive thymine analogs in the PAM-distal region. Distinct from other inducible Cas9 systems, vfCRISPR pre-binds to genomic DNA targets but remains catalytically inactive in the absence of light (Fig. 1A, top). The caging groups create steric hindrance that prevents hydrogen bonding between the modified nucleotides and target sequence, locking Cas9 protein in a cleavage-deficient conformation with partially unwound DNA duplex (Sternberg et al. 2015). Upon exposure to light, the caging groups are rapidly removed, allowing the pre-bound Cas9 to fully unwind target DNA and induce synchronous DSBs within seconds across all cells.

**Figure 1.**
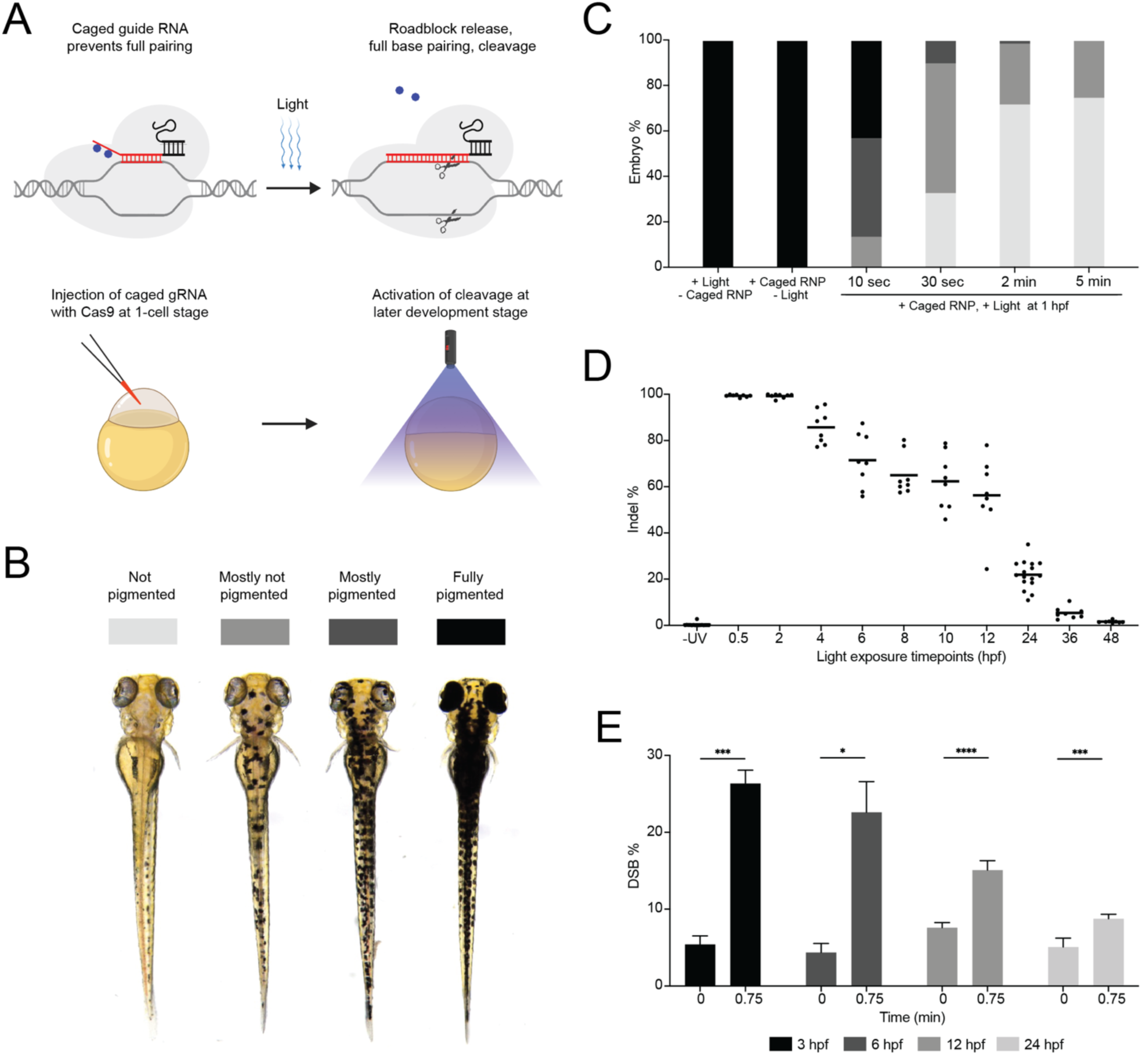
Light-activated CRISPR for rapid DSB induction in zebrafish embryos. **A.** Schematics of guide RNA uncaging with light to activate CRISPR-Cas9 genome editing (top) and implementation of light-activated CRISPR in the zebrafish embryo (bottom). Caging groups are shown in blue, base paired regions of the crRNA and target DNA are shown in red. **B.** Example images of four categories of melanin pigmentation loss after *tyr* gene disruption in 72 hours post-fertilization (hpf) zebrafish larvae. The associated shades of grey above each pigmentation loss category act as the legend for panels C & D. **C.** Quantification of pigmentation loss in uninjected embryos treated with 5 minutes of light at 1 hpf, embryos injected with caged *tyr* RNPs but not treated with light, and injected embryos exposed to various durations of light as indicated. All light exposures were conducted at 1 hpf. **D.** Amplicon sequencing analysis of *tyr* mutation frequency in caged *tyr* RNP-injected embryos not treated with light and injected embryos treated with 2 minutes of light (1-cell to 12 hpf) or 5 minutes of light (24 to 48 hpf). Indel percentage is defined as the number of reads containing insertions or deletions at the target site divided by the total number of reads. Each dot represents a single embryo. **E.** DSB frequency measured by DSB-ddPCR before light treatment and 0.75 minutes after light treatment in caged *tyr* RNP-injected 3, 6, 12, and 24 hpf embryos. Error bars represent standard deviation between 3 biological replicates for 3 and 6 hpf samples and 6 biological replicates for 12 and 24 hpf embryos. * = p ≤ 0.05, *** = p ≤ 0.001, **** = p ≤ 0.0001.

To test vfCRISPR activity in zebrafish, we designed a cgRNA targeting the *tyrosinase (tyr)* gene that encodes an enzyme responsible for the conversion of tyrosine to melanin in melanophores (Camp and Lardelli 2001). Zebrafish carrying null mutations in *tyr* lack melanin pigmentation, producing a phenotype that can be visually scored as a proxy for genome editing efficiency (Takasugi et al. 2022). We microinjected RNPs containing *tyr* cgRNA into single-cell zebrafish embryos and exposed them to various light dosages at one hour post-fertilization (hpf) (Fig. 1A, bottom, see Methods). At 72 hpf, embryos were examined and classified into one of four phenotypic categories ranging from zero to complete pigmentation loss (Fig. 1B). While treatment using light or RNP alone did not trigger any pigmentation loss, light-activation of RNPs in embryos induced pigmentation loss in a dosage-dependent manner (Fig. 1C). Importantly, 2 minutes of light exposure was sufficient to drive complete pigmentation loss in >70% of embryos without causing development defects (Fig. 1C, S1A), a result comparable to the phenotype penetrance using RNP with unmodified gRNA targeting the same site (Fig. S1B).

To determine whether vfCRISPR can be efficiently activated across embryo development, we treated embryos with light at timepoints from immediately after microinjection to 12 hpf (see Methods) and assessed pigmentation loss at 72 hpf. Light activation of vfCRISPR in the first two hours of embryo development produced complete pigmentation loss in >75% of embryos (Fig. S1C). vfCRISPR activity progressively declined in later stages of development; for example, light activation at 8 hpf resulted in only ∼20% of embryos exhibiting complete pigmentation loss. However, even after light activation at 12 hpf, nearly all embryos lost some pigmentation, suggesting that light-activated CRISPR activity persists long into embryo development.

The pigmentation loss assay lacks the sensitivity to detect edits at low efficiencies and does not report non-null mutations, since it requires both alleles of *tyr* to be inactivated in melanophores to result in a visible phenotype. Because of these issues, we quantified vfCRISPR activity directly by sequencing the targeted *tyr* locus from individual injected embryos after light activation at timepoints from immediately after injection to 48 hpf. Consistent with the phenotypic assay, <1% of alleles contained an insertion or deletion (indel) in injected embryos that were not exposed to light, while early activation (<2 hpf) led to nearly 100% mutated alleles (Fig. 1D). With this more sensitive and quantitative assay, we detected sustained and efficient editing with light activated vfCRISPR - ∼60% of *tyr* alleles contained indels after light activation between 6 and 12 hpf (Fig. 1D), consistent with the modest pigmentation loss at those stages. Notably, even at 24 hpf, light activation induced indels in ∼20% of alleles, demonstrating high vfCRISPR activity in a broad temporal window throughout zebrafish embryogenesis.

We hypothesized that vfCRISPR should instantaneously induce DSBs in embryos upon light stimulation, similar to its activity in cultured cells (Liu et al. 2020). To test this, we injected *tyr-*targeting RNPs into embryos and light activated them at 3, 6, 12, and 24 hpf. We then immediately collected genomic DNA from embryos for direct DSB quantification using digital droplet PCR (DSB-ddPCR) (Rose et al. 2017). In 3 and 6 hpf embryos, we found that ∼25% of target DNA was cleaved 45 seconds after exposure to light, indicating rapid and efficient cleavage at the *tyr* locus at both embryonic timepoints (Fig. 1E). We found that DSB induction also occurred within 45 seconds in 12 and 24 hpf embryos, albeit at lower rates consistent with the reduced editing efficiencies we previously observed. Similar phenotypic penetrance, indel percentage, and DSB induction kinetics were observed using a second cgRNA targeting *sox32* (Fig. S2). Together, our data demonstrates that vfCRISPR is an efficient tool for generating defined DSBs rapidly and across the first day of zebrafish embryo development.

### MMEJ drives rapid DSB repair in the zebrafish embryo

During early embryonic stages (0-5 hpf), zebrafish cells divide extremely fast, spending most time cycling between DNA synthesis (S phase) and mitosis (M phase) (Vastenhouw et al. 2019; Siefert et al. 2015). DNA damage-induced checkpoint activation is mostly absent at these stages, limiting time for repair when DSBs occur. How does the embryo avoid genome instability caused by DSBs during these rapid cell cycles? One possibility is that DSBs are repaired faster than cell division. Alternatively, DSB repair can be slow, but then cleaved chromosomes must be physically held together through a round of DNA replication and cell division before being repaired in daughter cells, as has been observed in cultured cells (Leimbacher et al. 2019; De Marco Zompit et al. 2022).

To distinguish between these possibilities, we measured the rate of indel formation in the zebrafish embryo using vfCRISPR and time-resolved amplicon sequencing. Single-cell embryos were injected with *tyr*-targeting RNPs and light activated at 3 hpf. Subsequently, genomic DNA from individual embryos was collected at a series of time points from 0.75 to 180 minutes after light activation and analyzed for indel presence using amplicon sequencing (Fig. 2A). Before DSB induction, less than 1% of alleles contained an indel, a value near the error rate of Illumina sequencing. After DSB induction, we observed ∼5% indels within 15 minutes, which rose to 12% indels within thirty minutes (Fig. 2B). Our data demonstrate that DSB repair is fast enough in the early zebrafish embryo to outpace the cell cycle, given that embryonic cell cycles last approximately 30-45 minutes in 3-4 hpf zebrafish embryos (Siefert et al. 2015; Dey et al. 2023).

**Figure 2.**
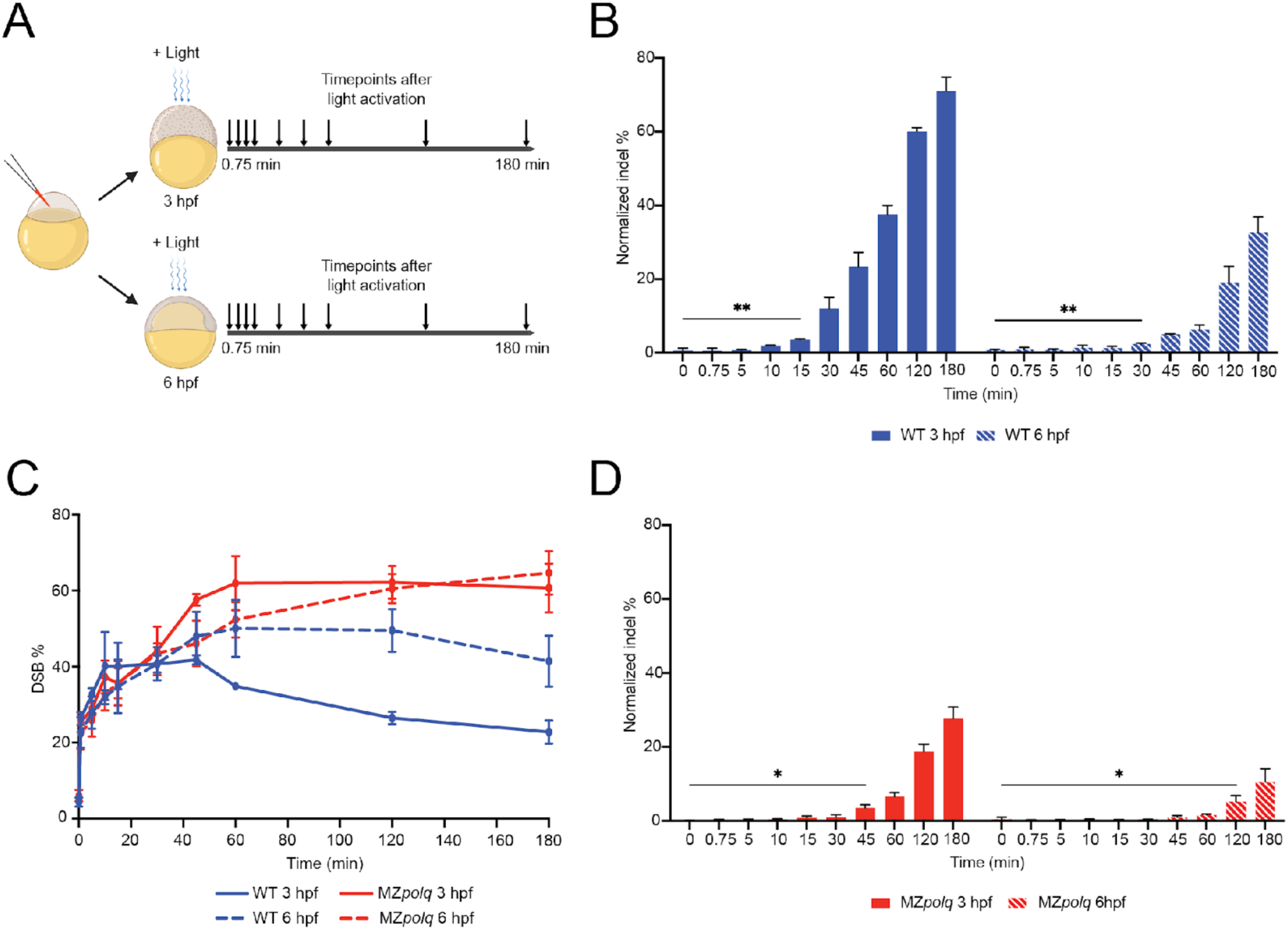
Fast DNA repair in early-stage embryos depends on MMEJ. **A.** Experimental design for collecting DNA from embryos across a time course after light-activated CRISPR cleavage. Time course was conducted after light activation at 3 hpf and 6 hpf. All embryos were injected with caged *tyr* RNP at the 1-cell stage. Each collection timepoint consists of ∼20-22 embryos. **B.** Normalized indel frequency before light treatment and at the indicated timepoints from 0.75 minutes to 180 minutes after light treatment in caged *tyr* RNP-injected 3 hpf and 6 hpf wildtype (WT) embryos. To account for the difference in DSB levels between WT embryos of different developmental stages, we calculated a normalized indel frequency instead of plotting raw indel frequency (see Methods). **C.** DSB frequency before light treatment and at timepoints from 0.75 minutes to 180 minutes after light treatment in caged *tyr* RNP-injected 3 hpf WT, 6 hpf WT, 3 hpf MZ*polq*, and 6 hpf MZ*polq* embryos. DSB frequency was quantified using ddPCR. **D.** Normalized indel frequency before light treatment and at timepoints from 0.75 minutes to 180 minutes after light treatment in caged *tyr* RNP-injected 3 hpf and 6 hpf MZ*polq* embryos. Error bars in all plots represent standard deviation between 3 biological replicates. * = p ≤ 0.05, ** = p ≤ 0.01.

When gastrulation initiates in the zebrafish embryo at ∼5 hpf, the cell division rate slows down dramatically and becomes asynchronous (2-3 hours per cycle) (Siefert et al. 2015; Dey et al. 2023). To test whether DSB repair shifts in response to these changes in cell cycle dynamics, we induced DSBs in embryos at 6 hpf and analyzed indel formation. We found that indels emerged far slower at the later stage, not until 30 minutes after DSB induction (Fig. 2B), even though the induced DSB levels were similar in embryos at 3 and 6 hpf (Fig. 1E). Because indels report only mutagenic repair events, we directly measured DSB and their resolution over time using DSB-ddPCR (Fig. 2C). Although the initial kinetics of DSB accumulation are similar, the number of alleles containing a DSB starts to decline within 60 minutes after CRISPR activation at 3 hpf (blue, solid line), but stays higher at 6 hpf (blue, dash line). These results suggest that DNA repair kinetics may decline alongside the slowing of the cell cycle as embryo development proceeds. DSB repair in the early-stage zebrafish embryo depends on the MMEJ pathway (Thyme and Schier 2016; Carrara et al. 2025). To test if rapid repair observed in 3 hpf embryos depends on MMEJ, we compared DSB perdurance between wild-type (WT) and homozygous maternal-zygotic *polq* mutant embryos (MZ*polq*, Fig. S3A). The *polq* gene encodes DNA polymerase theta, which catalyzes the templated DNA synthesis for MMEJ; MZ*polq* mutant embryos are deficient in MMEJ repair (Thyme and Schier 2016; Carrara et al. 2025). When light activated at 3 hpf, MZ*polq* embryos maintained high DSB levels for hours (red, solid line), sharply contrasting the rapid repair seen in WT embryos (blue, solid line) (Fig. 2C). At 6 hpf, DSB resolution was also slightly slower in MZ*polq* embryos, but DSB remained high in both WT and MZ*polq* embryos (blue and red dash lines), suggesting a reduced contribution of MMEJ to DNA repair at this stage. The lack of DSB resolution in MZ*polq* embryos suggested a delay in DNA repair, which should also impact indel formation. We found that the emergence of indels was ∼3 fold slower in 3 hpf and 6 hpf MZ*polq* embryos compared to WT embryos (Fig. 2D). Together, these results indicate that the rapid indel emergence in early embryos is largely driven by *polq*-mediated MMEJ.

### Mathematical modeling of DNA repair kinetics in the zebrafish embryo

The rate of indel accumulation that we measure with amplicon sequencing integrates both repair kinetics and repair fidelity. The slower rates of indel accumulation in 6 hpf WT embryos and in MZ*polq* embryos could be due to slower repair kinetics or higher fidelity repair, or both. To quantitatively understand DNA repair kinetics and fidelity in the zebrafish embryo, we adapted a coupled ODE model (Brinkman et al. 2018; Liu et al. 2020) that captures the dynamics of CRISPR-induced double-strand break (DSB) formation and repair. The model incorporates the rate of DNA cleavage (*k*_c_) and two distinct repair outcomes pathways: perfect repair (*k*_p_), in which the DNA sequence is restored without alteration and remains susceptible to recutting, and mutagenic repair (*k*_m_), in which indel formation disrupts the CRISPR target site and prevents further cleavage (Fig. 3A). We trained the model separately using 3 hpf and 6 hpf datasets to account for differences in Cas9 RNP concentration and the exponential increase in available target sites during rapid embryonic cell divisions. The model provided an excellent fit to the data, with high R^2^ values and low residuals across all conditions (Fig. 3B, S4, Table S2).

**Figure 3.**
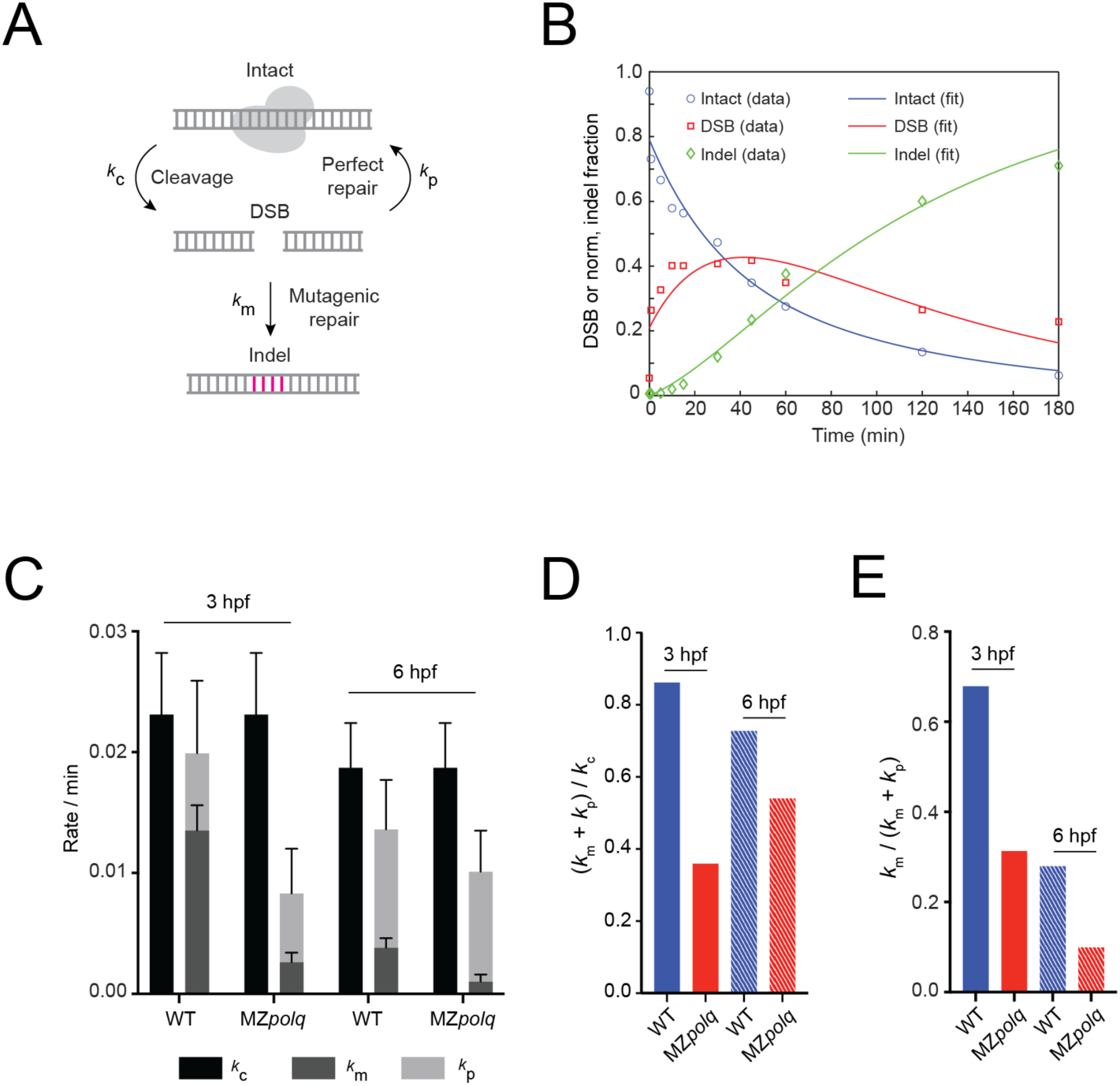
Kinetic modeling of Cas9 cleavage and repair kinetics in embryos. **A.** Schematic of the DSB repair model based on stochastic transitions between intact, DSB-containing, and indel states of target DNA. Cas9 induces a DSB, which is subsequently repaired, yielding an indel or restoring the original target sequence. *k*_*p*_and *k*_*m*_ are rate constants of perfect and mutagenic repair, respectively. *k*_*c*_is the cleavage rate constant for Cas9. **B.** Time course of DSB (red) and indel (green) fractions following light activation at 3 hpf in WT. The sum of DSB, normalized indel, and intact DNA equals 1. Normalized indel = measured indel × (1 – DSB). The solid lines indicate the best fit. R^2^ = 0.963. **C.** Estimated model parameters (*k*_*c*_, *k*_*p*_, *k*_*m*_) at 3 hpf and 6 hpf for WT and MZ*polq*, obtained by best model fitting. Error bars indicate 95% confidence intervals. **D.** DSB repair efficiency estimated by the fraction of total repair rate over cleavage rate at 3 hpf and 6 hpf for WT and MZ*polq*, defined as (*k*_*m*_ + *k*_*p*_) / *k*_*c*_. **E.** DNA mutation likelihood estimated by the fraction of mutagenic repair rate over total repair rate at 3 hpf and 6 hpf for WT and MZ*polq*, defined as *k*_*m*_ / (*k*_*m*_ + *k*_*p*_).

Analysis of the fitted parameters revealed three key insights. First, DNA repair occurs rapidly - substantially faster than would be inferred from indel accumulation alone due to the contribution of perfect repair - and far faster than the cell cycle itself (Fig. 3C). In 3 hpf WT embryos, the combined perfect and mutagenic repair rate closely matches the rate of DSB generation (Fig. 3D), indicating an efficient DSB detection and repair system despite the lack of cell cycle checkpoints at that stage. Second, disruption of MMEJ significantly reduced the rate of mutagenic repair with relatively little effect on perfect repair, demonstrating that MMEJ is a major contributor to mutagenic outcomes (Fig. 3C). Third, we infer the fraction of DSB undergoing mutagenic repair at 3 hpf (∼65%) is higher than at 6 hpf (∼30%), based on its contribution to the combined repair rates (Fig. 3E). Therefore, the observed decrease in indel rate in 6 hpf embryos is due to the combination of decreasing repair kinetics and increasing perfect repair rate. Together, these findings indicate that *polq*-mediated MMEJ is the dominant pathway driving rapid and mutagenic DSB repair in the early embryo, with its relative contribution diminishing during gastrulation as the cell cycle slows.

### The contribution of NHEJ to DNA repair progressively increases during development

The diminished role of MMEJ and increase of perfect repair in 6 hpf embryos suggests that other pathways begin to participate in DSB repair as embryo development proceeds. To assess the engagement of distinct repair pathways, we analyzed the repair products for mutational signatures after inducing targeted DSBs at the *tyr* or *sox32* loci at timepoints spanning early and later developmental stages. Because different repair pathways generate distinct outcomes, these signatures provide a quantitative readout of pathway usage. Indeed, a wide spectrum of indels were observed ranging from single-nucleotide insertions to larger deletions up to 25 bp (Fig. S5, S6B, S7B).

To simplify our analysis, we classified ‘+1 bp insertions’ and ‘short deletions (1-4 bp)’ as signatures of NHEJ, and ‘templated insertions’ together with ‘microhomology-mediated larger deletions (5-10 bp)’ as signatures of MMEJ, following precedents from the literature (Hussain et al. 2021; Schimmel et al. 2019; Sishc and Davis 2017; Chandramouly et al. 2021). Combined and individual NHEJ and MMEJ signatures were plotted to quantify the change of pathway usage over time (Fig. 4A, Fig. S6, Fig. S7). At the *tyr* locus, MMEJ accounted for ∼ 60% of indel when DSBs were induced at 2 hpf, whereas NHEJ contributed only ∼6%, again supporting that MMEJ is the dominant repair pathway at early stages. As development progressed, NHEJ activity increased steadily by five fold to ∼30% by 12 hpf, meanwhile MMEJ activity decreased to just below 40%. A similar trend was observed at the sox32 locus (Fig. S7). Our results reveal a gradual shift from MMEJ to NHEJ pathway usage in DNA repair during zebrafish embryogenesis.

**Figure 4.**
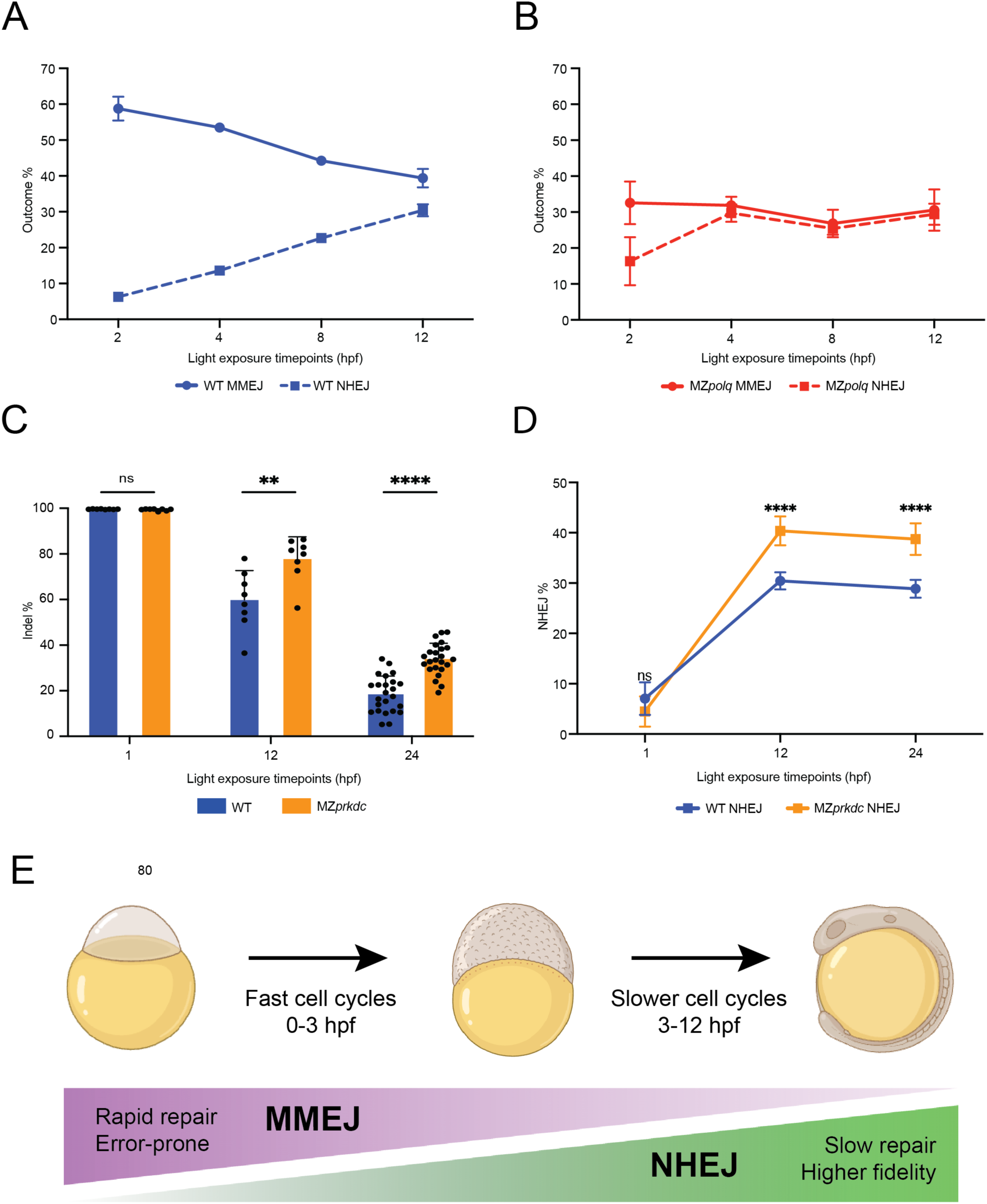
A shift in repair outcomes and fidelity as embryogenesis proceeds. **A.** Mutation frequency of MMEJ-associated repair outcomes (templated insertions and microhomology-mediated 5–10 bp deletions) and mutation frequency of NHEJ-associated repair outcomes (1 bp insertions and 1-4 bp deletions) at the *tyr* locus in caged *tyr* RNP-injected WT embryos after light activation at the indicated time points. Individual embryos were collected at 26–28 hpf for amplicon sequencing. Error bars represent standard deviation between 8 biological replicates. **B.** Mutation frequency of MMEJ-associated repair outcomes (templated insertions and microhomology-mediated 5–10 bp deletions) and mutation frequency of NHEJ-associated repair outcomes (1 bp insertions and 1-4 bp deletions) at the *tyr* locus in caged *tyr* RNP-injected MZ*polq* embryos after light activation at the indicated time points. **C.** Indel frequency at the *tyr* locus in caged *tyr* RNP-injected WT and MZ*prkdc* embryos after light activation at 1, 12, or 24 hpf. Embryos after light activation at 1 and 12 hpf were collected at 26–28 hpf and embryos after light activation at 24 hpf were collected at 48 hpf for amplicon sequencing. Each dot represents a single embryo. Error bars represent standard deviation between 8 biological replicates for 1 and 12 hpf embryos and 24 biological replicates for 24 hpf embryos. **D.** Mutation frequency of NHEJ-associated repair outcomes at the *tyr* locus in caged *tyr* RNP-injected WT and MZ*prkdc* embryos after light activation at 1, 12, or 24 hpf. ** = p ≤ 0.01, **** = p ≤ 0.0001, ns = not statistically significant. **E.** Proposed model for DNA repair pathway usage during zebrafish embryo development. The error-prone MMEJ pathway (purple gradient) dominates during the early embryo stage when the cell cycle is fast. The higher-fidelity NHEJ (green gradient) takes over at later embryo development when the cell cycle becomes slower.

Previous studies have linked minimal NHEJ activity observed in early embryos to the low expression of core NHEJ factors (Thyme and Schier 2016; Carrara et al. 2025). Alternatively, MMEJ may dominate repair pathway choice through competition at a DSB, even when NHEJ components are present and active. To test for potential competition between MMEJ and NHEJ, we repeated the time-course experiment in MZ*polq* mutant embryos. In contrast to WT embryos, the gradual increase in NHEJ signatures and concomitant decrease in MMEJ signatures were largely abolished in MZ*polq* mutants (Fig. 4B). Notably, NHEJ activity reached ∼16% at 2 hpf in mutant embryos, compared with ∼ 6% in WT embryos, and further increased to ∼30% by 4 hpf, a level that was not observed until 12 hpf in WT embryos. While these quantitative differences are caveated by the general delays and failures of DNA repair in MZ*polq* embryos, they still suggest that NHEJ is already active during early embryo stages but is outcompeted by *polq*-mediated MMEJ. Conversely, MMEJ associated signatures were reduced to ∼30% at 2 hpf in MZ*polq* mutant embryos and remained unchanged over the course of the experiment, suggesting the presence of *polq*-independent MMEJ mechanisms during zebrafish embryogenesis.

Although NHEJ is thought of as an error-prone repair pathway, it can mediate DSB repair with up to 90% fidelity after Cas9 cleavage (Liu et al. 2020). We therefore hypothesize that the transition toward NHEJ provides a more faithful route than MMEJ for DSB repair after embryo gastrulation. To test this, indel frequency was measured in WT embryos after DSB induction at 1, 12, and 24 hpf. Nearly 100% indel was observed at the *tyr* locus when DSB was induced 1 hpf and substantially less indels were detected at both 12 and 24 hpf (Fig. 4C).

The reduction in indels at 12 and 24 hpf could be consistent with higher-fidelity repair when NHEJ is engaged, but are certainly also influenced by reduced Cas9 activity due to dilution or degradation of vfCRISPR RNPs during development. To distinguish between these possibilities, we disrupted NHEJ fidelity by knocking out the *prkdc* gene (Fig. S3B). *prkdc* encodes DNA-PKcs, which is recruited to a DSB to protect DNA ends from excessive resection and coordinate a biased repair towards NHEJ (Yue et al. 2020). *prkdc* is required for V(D)J recombination in zebrafish and other vertebrates (Moore et al. 2016; Mashimo et al. 2012). While *prkdc* mutant zebrafish lack a functional adaptive immune system, they can be raised in the lab setting to fertile adulthood (Moore et al. 2016; Jung et al. 2016), unlike zebrafish deficient in other NHEJ components (Carrara et al. 2025).

We compared indel frequencies between WT and MZ*prkdc* mutant embryos. No difference was observed between the genotypes at 1 hpf (Fig. 4C), when NHEJ contribution to DNA repair is minimal (Fig. 4A,D). In contrast, after light activation at 12 or 24 hpf, MZ*prkdc* mutants had significantly increased mutation frequency when compared to WT controls. At 24 hpf, MZ*prkdc* embryos accumulated nearly two-fold higher indels, from 18% to 34%. This overall increase in indel frequency was driven by an increase in NHEJ signatures in MZ*prkdc* embryos after DSB induction at 12 or 24 hpf (Fig. 4D, S8). Together, these data show that DNA-PKcs is critical for faithful repair of DSBs at later stages of zebrafish embryo development, consistent with our kinetic modeling results (Fig. 3E).

At the *sox32* locus, we observed a similar pattern - repair outcomes were similar after CRISPR activation at 1 hpf, but NHEJ outcomes had significant changes in their likelihood after CRISPR activation at 12 and 24 hpf (Fig. S9). We noticed some distinctions between the repair outcomes at these two loci - notably, a difference in the likelihood of 1 bp insertions, which were ∼5- to 10-fold more likely at the *tyr* locus. We suspect that sequence and loci dependent effects on DNA repair may explain these differences, as observed in other systems (Allen et al. 2018). Regardless of locus-specific repair outcomes, we found that disruption of NHEJ had a strong effect on DNA repair at later stages of zebrafish embryogenesis (Fig. S8, S9). Together, these data support a model in which a shift from rapid and mutagenic MMEJ to slower and more faithful NHEJ repair coincides with the slowing of the embryonic cell cycle during gastrulation (Fig. 4E).

## DISCUSSION

In this study, we adapted a light-activated vfCRISPR system to enable precise temporal control of DSB generation in zebrafish embryos. A Cas9-cgRNA complex “parked” at its target locus can be activated with a brief pulse of light to induce DSBs within seconds and produce mutations with close to 100% efficiency (Fig. 1). By synchronizing DSB generation, we directly measured the pace of DNA repair in a vertebrate embryo for the first time. We found that DSBs in the early-stage zebrafish embryo are detected and repaired with remarkable speed - mutations emerged at target sites within minutes of CRISPR activation (Fig. 2B). To our knowledge, this is the fastest repair of DSBs observed in any living cell.

The choice of DSB repair pathway has been a long-standing mystery (Chang et al. 2017; Ceccaldi et al. 2016; Shrivastav et al. 2008; Kumari et al. 2025). In mammalian cells, NHEJ is the predominant repair pathway in G1, with HR playing an increasingly prominent role during S and G2 phases. Given its highly mutagenic nature, MMEJ has been proposed as a backup repair mechanism when NHEJ and HR are compromised or unavailable (Brambati et al. 2023; Martin et al. 2025; Sfeir and Symington 2015). Surprisingly, our data show that zebrafish embryonic cells instead depend on the MMEJ machinery to drive DSB repair, with NHEJ serving as a secondary pathway. At early developmental stages (2-3 hpf), rapid DSB repair is largely dependent on *polq*-mediated MMEJ (Fig. 2C), and MMEJ outcomes are far more frequent than NHEJ outcomes (Fig. 4A). In MZ*polq* mutant embryos, NHEJ activity only modestly compensates for repair outcomes (Fig. 4B); however, DSB resolution is markedly slower under these conditions (Fig. 2C and Fig 3C). Our findings extend previous work establishing that *polq*-mediated MMEJ is essential for early embryonic survival in zebrafish after CRISPR-induced DSBs (Thyme and Schier 2016; Carrara et al. 2025).

Previous studies have suggested distinct DSB repair pathway usage during zebrafish embryogenesis. Embryos lacking *polq* are initially sensitive to ionizing radiation, which induces a mixture of DNA lesions, but start to acquire resistance at 6 hpf and become as radiation-tolerant as WT embryos at 24 hpf (Thyme and Schier 2016; Bladen et al. 2007; Carrara et al. 2025). In contrast, knockdown of key NHEJ components renders embryos sensitive to irradiation at 6 hpf (Bladen et al. 2005, 2007). Using light-activated CRISPR and targeted sequencing, we unambiguously show that DSB repair pathway usage shifts from MMEJ to NHEJ as development progresses. MMEJ is the primary mechanism driving rapid DSB repair at early stages (Fig. 2 and 3), whereas NHEJ, although present, is not actively engaged. The pace of DSB repair declines progressively with embryo development, coinciding with a change in repair outcomes from MMEJ-signature mutations to NHEJ signatures (Fig. 4). Finally, MZ*prkdc* mutant embryos with disrupted NHEJ have similar repair outcomes to WT embryos at 1 hpf, but significantly altered outcomes at 12 or 24 hpf (Fig. 4). Together, these findings support a model in which DNA repair shifts from MMEJ to NHEJ during the first 12 hours of zebrafish development.

So why is MMEJ the dominant pathway for DSB repair in early zebrafish development? We propose the following explanation (Fig. 4E). In early-stage zebrafish embryos, the cell cycle proceeds at a relentless pace (every 15 minutes) without cell cycle checkpoints. The priority for DSB repair is speed at the expense of fidelity, preventing chromosome mis-segregation or catastrophic loss of fragments that could happen at DNA replication. Our data show that MMEJ repair in the early-stage embryo can occur within minutes after a DSB (Fig 2B and C), but with low fidelity - 65% of DSBs yield a mutation (Fig. 3E). As the pace of cell cycles slows a few hours after fertilization, and cell cycle checkpoints emerge, the embryo has bought the time to engage slower but more accurate repair mechanisms such as NHEJ, which yield a perfect repair outcome ∼70% of the time. There is enormous variation in the pace of development across the tree of life (Ebisuya and Briscoe 2018; O’Farrell et al. 2004). Applying light-activated CRISPR in other organisms may uncover the evolutionary significance behind the correlations we observe between developmental tempo and DNA repair pathway choice.

Several unexpected findings from our study suggest avenues for future research. First, it remains unclear why MMEJ proceeds more rapidly than NHEJ. Recent work has shown that NHEJ requires the stepwise assembly and disassembly of a complex core repair machinery, taking ∼25 minutes until DSB resolution (Mikhova et al. 2024). In contrast, MMEJ relies on fewer components and may therefore be intrinsically less time-consuming. Direct measurements of MMEJ kinetics using high resolution imaging or time-resolved ChIP-seq may help test this possibility (Liu et al. 2020; Zou et al. 2022; He et al. 2024; Marin-Gonzalez et al. 2025). Second, the mechanisms coordinating the handoff from MMEJ to NHEJ and the extent to which these pathways may intersect remains poorly understood. mRNAs encoding the components of both pathways are present throughout embryogenesis (Carrara et al. 2025; Thyme and Schier 2016; Sur et al. 2023). However, mRNA levels may not reflect the actual protein abundance, stoichiometry, and availability of DNA repair complexes in the embryo. Third, some repair products consistent with MMEJ, particularly deletions larger than 10 bp, persist even in the absence of *polq* (Fig. S6C, S7C). We suspect that *polλ* or other replication-associated polymerases can mediate *polq*-independent MMEJ to yield these mutations (Chandramouly et al. 2023; Feldman et al. 2021; Carvajal-Garcia et al. 2020). It would be interesting to test whether these polymerases are recruited with slower kinetics than *polq*, potentially explaining the delayed DSB resolution observed in MZ*polq* mutant embryos. Alternatively, *polq* may engage shorter or simpler microhomologies that enable faster initiation of DNA synthesis and DSB repair.

Finally, vfCRISPR technology itself could be further improved. In our study, cgRNAs remain active after light exposure, allowing repeated cycles of DNA damage and repair until a substantial indel is generated. Incorporating photocleavable guide RNAs to rapidly deactivate Cas9 could control its activity to one round of DSB generation and repair (Zou et al. 2021), simplifying the interpretation of repair fidelity. In addition, the cgRNAs we used target a single genomic locus, limiting the generalizability of our conclusions across the complex genome. Recently, high-throughput cgRNAs capable of simultaneously targeting more than 100 sites have been engineered to study DSB repair in human cells (Zou et al. 2022; Marin-Gonzalez et al. 2025). Adapting these technological advances would enable genome-wide interrogation of DSB repair dynamics across embryogenesis.

## MATERIALS AND METHODS

### Zebrafish husbandry

All zebrafish work was performed at the CBRZ facility of the University of Utah. This study was approved by the Office of Institutional Animal Care & Use Committee at the University of Utah (Protocol 00001701).

### Generation of mutant lines

We mutated *polq* and *prkdc* using CRISPR-Cas9 (Figure S3). Two SpCas9 crRNAs that target exon 3 of *polq* and two cRNAs that target exons 6 and 7 of *prkdc* were ordered from IDT. We assembled dgRNAs from crRNAs and tracRNA following instructions from the manufacturer (IDT). We assembled an injection mix using this recipe - 1.5 μL dgRNAs at equimolar ratio (∼800 ng/uL), 1.5 μL 1M KCl (Sigma-Aldrich), 1 μL SpCas9 protein (Alt-R® S.p. Cas9 nuclease, v.3, IDT) and 1 μL 1% phenol red (RICCA), assembled in that order. This mix was briefly vortexed and centrifuged to bring the solution to the bottom of the tube and placed on ice. We injected 1 nl of this mix directly into the cell of wild-type TU zebrafish zygotes. Injected embryos were raised to adulthood and incrossed to generate trans-heterozygote lines or outcrossed to wild-type to generate heterozygote lines. We identified F1 individuals with edited alleles containing large deletions using PCR with primers that flank the region targeted for editing.

For *polq,* trans-heterozygous mutants carrying two distinct large deletions were used to generate embryos for all experiments studying repair kinetics and outcomes. We mapped four specific alleles in these individuals with 504, 508, 509, and 512 bp deletions, all of which are predicted null alleles. Subsequently, we identified heterozygous adult fish with a 512 bp deletion which we used to establish homozygous mutant lines. Comparisons between trans-heterozygous and homozygous parents (carrying different deletion alleles) yielded the same consequences for DNA repair in the 3 hpf embryo. For *prkdc,* trans-heterozygous mutants carrying 230 bp and 233 bp deletions were used for all experiments. Maternal-zygotic *polq* (MZ*polq*) and *prkdc* (MZ*prkdc*) mutant embryos lack maternally contributed wild-type gene products and cannot zygotically produce wild-type gene products. MZ*polq* and MZ*prkdc* mutant embryos were generated by crossing mutant parents to each other.

The sequences of all guide RNAs, primers, and mutant alleles can be found in **Table S1**.

### Preparation of caged guide RNAs

crRNAs containing caged nucleotides (two or three light-sensitive, 6-nitropiperonyloxymethyl-modified deoxynucleotide caged thymine nucleotides that replace uracils, see Table S1 for details) were purchased from Biosynthesis Inc. Caged crRNAs and standard SpCas9 tracrRNAs (IDT) were reconstituted at 100 μM in duplex buffer (IDT). To prepare the crRNA:tracrRNA duplex, equal volumes of 100 μM caged crRNA and 100 μM tracrRNA were mixed, diluted to 25 μM with additional duplex buffer, and annealed by heating at 95 °C for 5 minutes followed by cooling to room temperature. The 25 μM caged crRNA:tracrRNA duplex was aliquoted in single-use tubes and stored at −20 °C.

### Assembling and injecting CRISPR-Cas injection mixes with caged guide RNAs

Microinjection mixes containing caged guide RNAs were assembled in 1.5 ml tubes in the following order: 1.5 µl of 25 μM crRNA:tracrRNA duplex, 1 µl of 1 M KCl (Sigma-Aldrich), 1.5 µl of SpCas9 protein (Alt-R® S.p. Cas9 nuclease, v.3, IDT) and 1 µl of 1% phenol red (RICCA). This mix was briefly vortexed and centrifuged to bring the solution to the bottom of the tube and placed on ice. We injected 1.5 nl of this mix directly into the cell of zebrafish zygotes.

### Light activation of CRISPR-Cas9 DNA cleavage

Embryos were raised at 28.5 °C after injection. To activate CRISPR-Cas9 mediated DNA cleavage, injected embryos were transferred into a single well of a 12-well tissue culture plate (VWR, Cat.# 10062-894) containing just enough E3 buffer to fully cover them and exposed to 365 nm light (1.3 J/cm²) by placing a flashlight (Jaxman, U1) directly on top of the transparent plastic lid. For embryos at developmental time points >8 hpf, the lid was removed to minimize UV attenuation. Exposure time was recorded starting from activation of the flashlight with a timer. All exposures were conducted in a windowless room to prevent incidental exposure to sunlight. Following exposure, embryos were transferred into Petri dishes filled with 28.5 °C E3 buffer and returned to a dark incubator.

### Pigmentation scoring

At 1 dpf, we screened to remove unfertilized or dead embryos. In most conditions, <10% of embryos exhibited toxicity, which we attribute mostly to injection artifacts. At 3 dpf, embryos were scored for pigmentation loss into one of four categories: fully pigmented (100% pigmentation), mostly pigmented (51–99% pigmentation), mostly not pigmented (6–50% pigmentation), and not pigmented (0–5% pigmentation).

### Preparing samples for CRISPR-Cas9 kinetics measurements

Embryos were harvested at different time points by incubating them in an 80 °C heat block for 10 minutes to inactivate the Cas9 RNP. Samples were subsequently cooled to room temperature, transferred to ice, and processed for genomic DNA extraction using the DNeasy Blood & Tissue Kit (Qiagen) according to the manufacturer’s instructions.

### Measuring DSBs with digital droplet PCR

We calculated the percentage of alleles containing a DSB using DSB-ddPCR <u>(Rose et al. 2017)</u>. Briefly, two amplicons were designed for each region of interest. The first amplicon (Target 1) includes the CRISPR target site, while the other amplicon (Target 2) is not targeted for cleavage and is used to normalize the DNA copy measurements. One dual-quenched qPCR probe (IDT) was designed for each amplicon. Primers and probes were designed using guidelines by Bio-Rad.

To set up ddPCR reactions to measure DSBs, 25 ng of purified genomic DNA was added to a 20 μL reaction containing the ddPCR Supermix for Probes (no dUTP) (Bio-Rad). This reaction has a final concentration of 900 nM of primers and 250 nM of probes. Droplets were created and DNA partitioned using Droplet Generation Oil for Probes, DG8 Gaskets, DG8 Cartridges, and the QX200 Droplet Generator (Bio-Rad). Droplets were subsequently transferred to a 96-well PCR plate and heat-sealed using PX1 PCR Plate Sealer (Bio-Rad). This plate was placed in a deep well thermocycler for PCR amplification with the following conditions: 95 °C for 10 min, followed by 40 cycles of (94 °C for 30 sec, 54.5 °C for 30 sec), 98 °C for 10 min and a final hold at 12 °C. After PCR, droplets are analyzed using the QX200 Droplet Reader (Bio-Rad). Droplets are distinguished as positive or negative in either fluorescent channel (FAM/HEX) based on whether the amplitude of fluorescence passes the global threshold set.

Using these raw data, DSB frequency was calculated using the following formula:

[Target 1(−), Target 2(+)] / ([Target 1(−), Target 2(+)] + [Target 1(+), Target 2(+)])

Target 1(−) indicates negative droplets for the Target 1 amplicon and Target 2(+) indicates positive droplets for the Target 2 amplicon.

### Sample preparation for amplicon sequencing

Genomic DNA was extracted from individual embryos at 1 dpf or 2 dpf using the following protocol. Individual embryos were incubated in 24 ul (1 dpf embryo) or 30 ul (2 dpf embryo) of alkaline lysis solution (25mM NaOH, 0.2 mM EDTA) at 95 °C for 30 minutes. After cooling to 4 °C, an equal volume of neutralizing solution (40 mM Tris HCl) was added. Genomic DNA was stored at 4 °C.

Oligo sequences for library preparation are in **Table S1**. Genomic DNA samples were amplified with PCR using Q5 Hot Start High-Fidelity 2X Master Mix (New England BioLabs M0494). PCR amplification as performed with the following conditions: 98 °C for 30 seconds, 25 cycles (*tyr),* 26 cycles (*sox32*) of {98 °C for 10 seconds, 65 °C for 10 seconds, 72 °C for 12 seconds}, 72 °C for 5 minutes, and 10 °C hold. PCR cleanup was performed using 1.2x Mag-bind TotalPure NGS (Omega BIO-TEK) following the manufacturer’s instructions. Dual-indexing PCR to uniquely index each sample was performed using Q5 Hot Start High-Fidelity 2X Master Mix with the following conditions: 98 °C for 30 seconds, 9 cycles of {98 °C for 10 seconds, 68 °C for 10 seconds, 72 °C for 15 seconds}, 72 °C for 5 minutes, and 10 °C hold. PCR cleanup was performed using 1x Mag-bind TotalPure NGS.

### Determination of somatic mutagenesis alleles

Samples were pooled at equimolar ratios, diluted, and sequenced using a MiSeq (Illumina) with MiSeq Reagent Kit v2 (300-cycles), as previously described (Gagnon et al., 2014). Collected data were analyzed using SIQ (Schendel et al., 2022) to identify and classify repair outcomes.

Templated insertions and deletions equal or larger than 5 bp with microbiology were classified as MMEJ outcomes <u>(Schimmel et al. 2019; Chandramouly et al. 2021)</u>. 1 bp insertions and deletions equal or smaller than 4 bp were classified as NHEJ outcomes <u>(Hussain et al. 2021; Sishc and</u> <u>Davis 2017)</u>.

### Normalized indel calculation from raw indel % and DSB %

We normalized indel % to factor in cleaved DNA that cannot be PCR amplified for indel measurements. Raw indel % is obtained from deep amplicon sequencing. DSB % is obtained from ddPCR. We define normalized indel % to be the indel percentage from all DNA at a particular locus, not just from PCR-amplifiable DNA. Normalized indel % = indel % × (100 – DSB %) / 100. The sum of DSB %, normalized indel %, and intact % equals 100%.

### Mathematical modeling of Cas9-induced DSB and indel formation

To quantify DNA repair kinetics in zebrafish embryos, we adapted a previously described model of DSB and indel formation <u>(Brinkman et al. 2018; Liu et al. 2020)</u>. Upon light stimulation (*t* = 0), a fraction of intact DNA (*B*_0_) is rapidly cleaved by pre-bound Cas9, generating DSBs. These breaks can subsequently undergo either perfect repair, restoring the original DNA sequence, or mutagenic repair, resulting in indel formation that prevents further Cas9 binding and cleavage. We define the rates of perfect and mutagenic repair as *k*_p_ and *k*_m_, respectively. We further define *k*_c_ as the apparent Cas9 cleavage rate, reflecting the combined contribution of two processes: (1) cleavage of initially unbound DNA targets by Cas9 molecules after light activation, and (2) re-cleavage of perfectly repaired DNA targets. Our model does not distinguish between these two contributions.

Based on these definitions, our model can be described as follows:

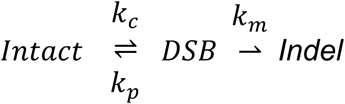

This reaction is further modeled as a set of coupled ordinary differential equations for WT and MZ*polq* conditions, respectively:

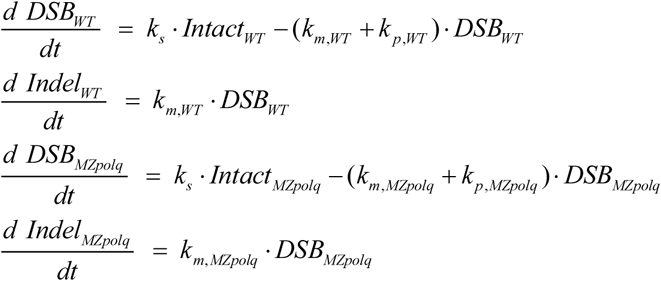

both datasets satisfying the normalization condition:

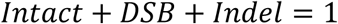

and initial condition:

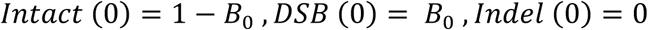

We simultaneously fit data acquired under WT and MZ*polq* conditions within a single optimization framework for each timepoint. Cas9 cleavage rate, *k*_c_, is constrained to be identical across both datasets for each timepoint because it should be independent of the presence of *polq.* Model parameters (*k*_*c*_, *k*_*p*_, *k*_*m*_) were fit to experimental data using the nonlinear least-square fitting function (lsqcurvefit) in MATLAB R2020a (MathWorks).

## ACKNOWLEDGEMENTS

We thank all members of the Gagnon and Liu laboratories, Jeffrey Farrell, David Grunwald, Nathan Lord, and Kazuyuki Hoshijima for helpful discussions and comments. We thank the Centralized Zebrafish Animal Resource (CZAR) and CBRZ staff, and the Genomics and DNA Peptide core facilities at the University of Utah. This project was supported by National Institutes of Health grants R35GM142950 (J.A.G.), R24OD035409 (J.A.G.), and R35GM150941 (Y.L.), and startup funds from the University of Utah (Y.L.). Y.L. is an inventor on a pending patent application covering the vfCRISPR technologies used in this study. The other authors declare no competing interests.

## SUPPLEMENTARY MATERIALS

Figs. S1 to S9

Tables S1 and S2

## Supplementary Materials

**Figure S1.**
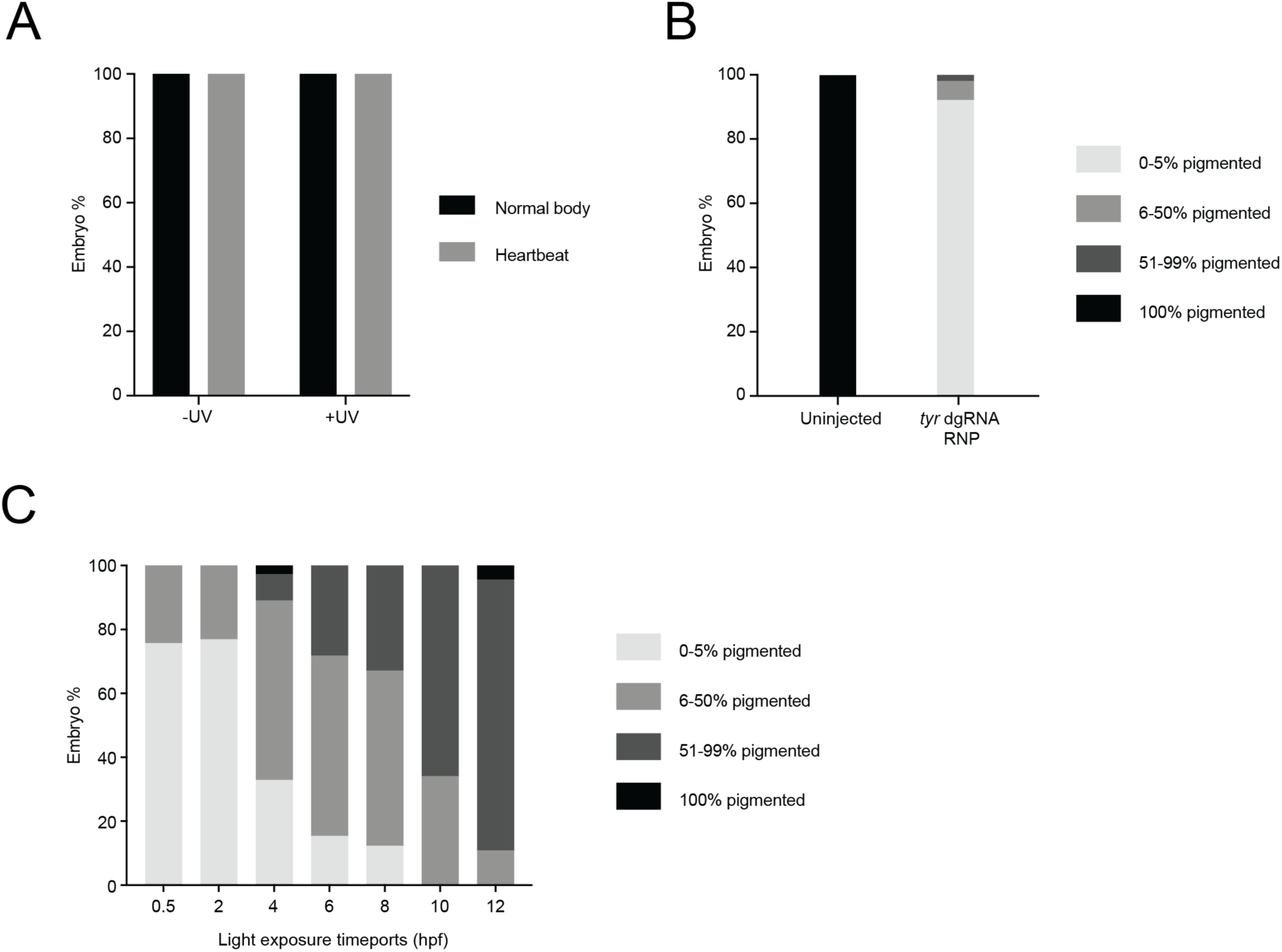
Control experiments for light-activated CRISPR. **A.** Quantification of normal body morphology and presence of a heartbeat in uninjected, untreated wild-type embryos or uninjected embryos treated with 2 minutes of light at 1 hpf and scored for each phenotype at 48 hpf. All embryos had normal body morphology and a heartbeat, indicating that this treatment of light is non-toxic to development. **B.** Quantification of pigmentation loss in uninjected embryos and embryos injected at the 1-cell stage with *tyr* dgRNA (constitutively active, not caged). Pigmentation categories are described in Figure 1B. **C.** Quantification of pigmentation loss in embryos injected with caged *tyr* RNPs and treated with 2 minutes of light at different times of development as indicated. Pigmentation categories are described in Figure 1B.

**Figure S2.**
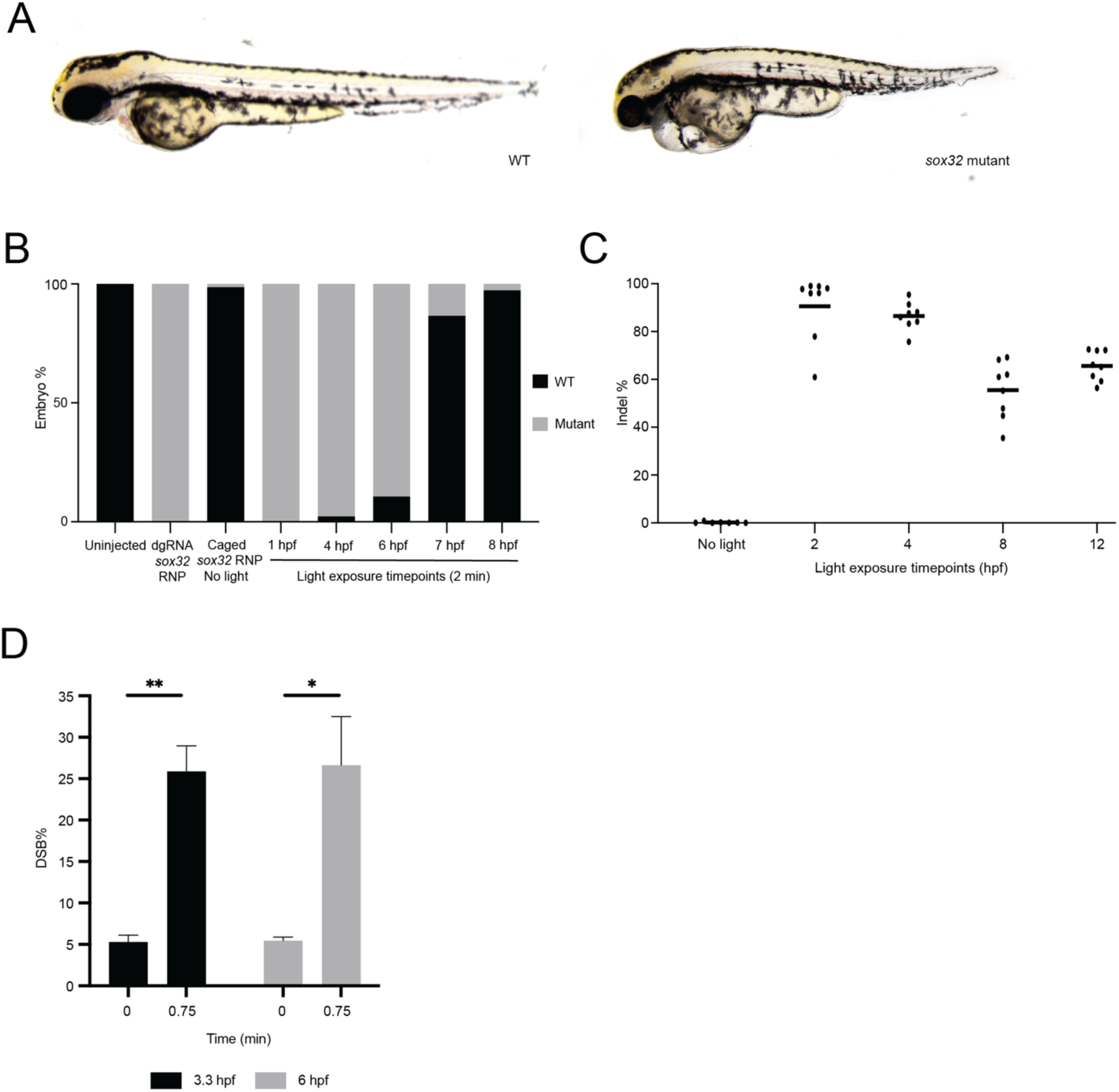
Temporal control of *sox32* editing in zebrafish embryos with light-activated CRISPR. A. Example images of wild-type (WT) or *sox32* mutant phenotypes in 48 hours post-fertilization (hpf) embryos. Key aspects of the phenotype are the shortened body length and heart edema. B. Quantification of *sox32* phenotype in uninjected embryos, embryos injected with *sox32* dgRNP (constitutively active, not caged), embryos injected with caged *sox32* RNP without light treatment, and embryos injected with caged *sox32* RNP and treated with 2 minutes of light at different times of development as indicated. C. Amplicon sequencing analysis of *sox32* mutation frequency in caged *sox32* RNP-injected embryos not treated with light, and injected embryos treated with 2 minutes of light at different times of development as indicated. Indel percentage is defined as the number of reads containing insertions or deletions at the target site divided by the total number of reads. Each dot represents a single embryo. D. DSB frequency before light treatment and 0.75 minutes after light treatment in caged *sox32* RNP-injected 3.3 and 6 hpf embryos. * = p ≤ 0.05, ** = p ≤ 0.01.

**Figure S3.**
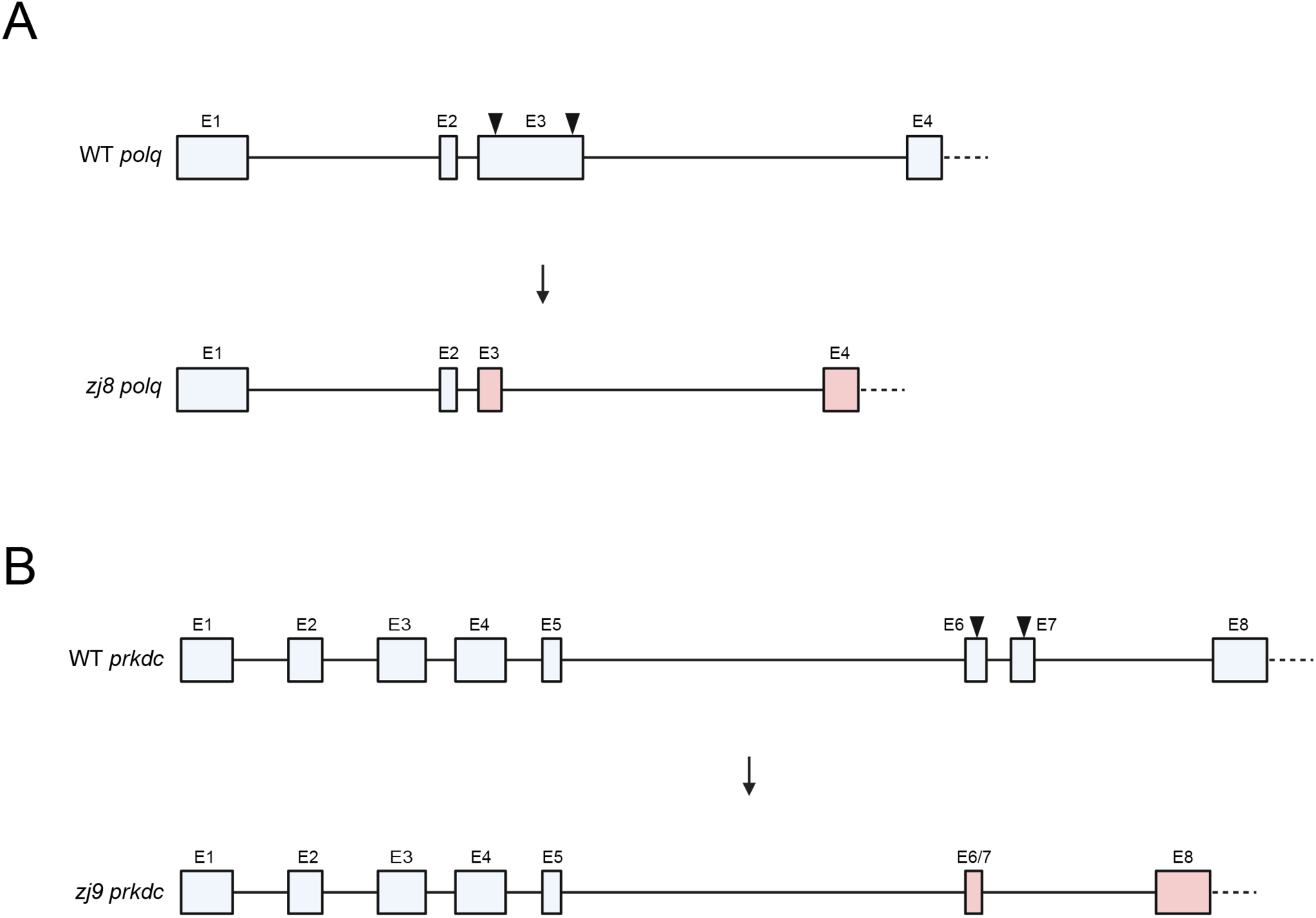
Diagrams of CRISPR mutants that disrupt the *polq* and *prkdc* genes. **A.** Design for mutating the *polq* gene, with flanking guide RNAs shown as arrowheads (above) and the *polq* allele with a large deletion that removes part of exon 3 (see Table S1 for sequences). **B.** Design for mutating the *prkdc* gene, with flanking guide RNAs shown as arrowheads (above) and the *prkdc* allele with a large deletion that removes part of exon 6 and exon 7 (see Table S1 for sequences).

**Figure S4.**
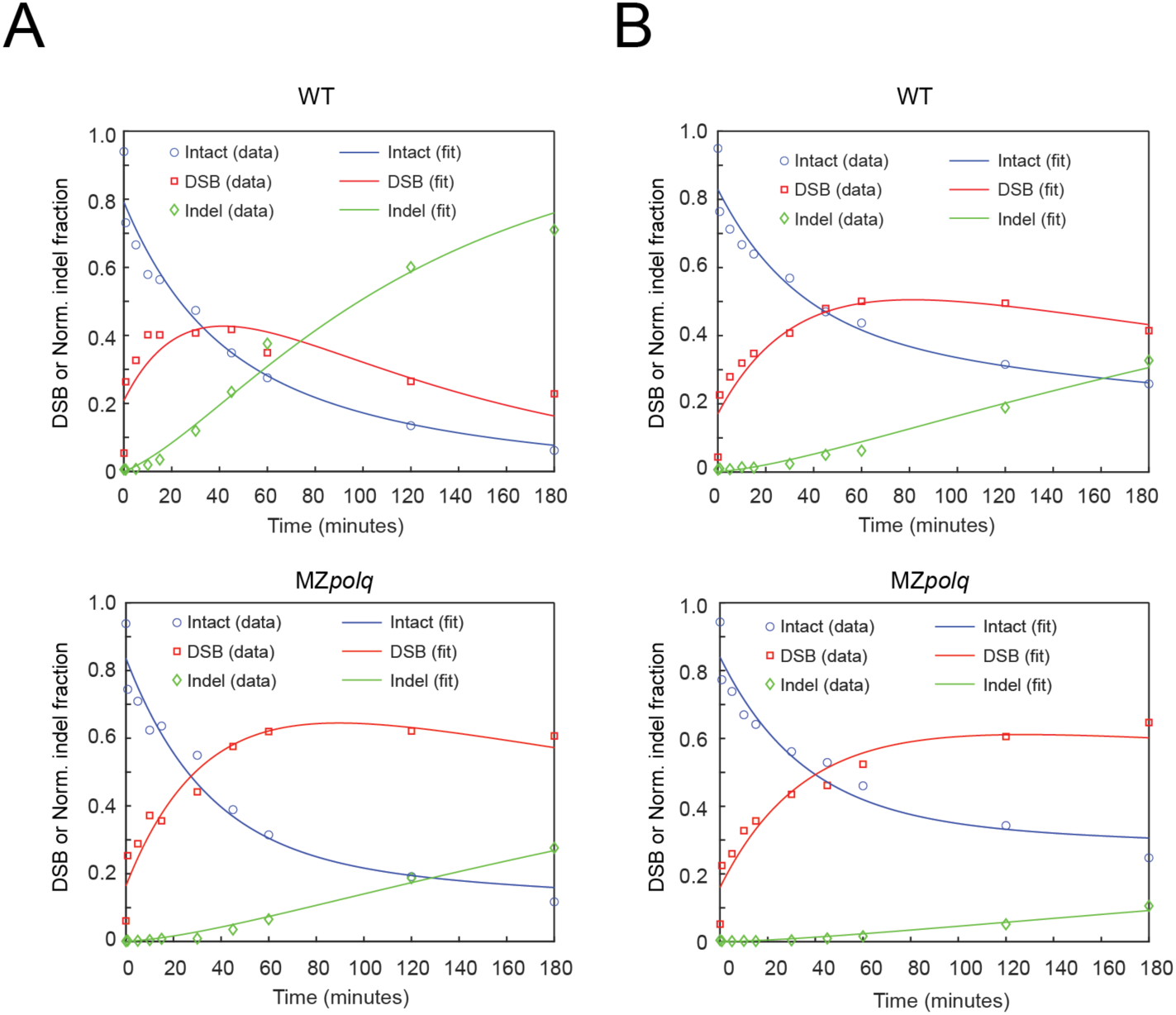
Kinetic modeling of Cas9-induced DSB induction and repair. **A.** Representative time courses of the intact (blue), DSB (red), and indel (green) fractions following light activation at 3 hpf in WT and MZ*polq* embryos. **B.** Representative time courses of the intact (blue), DSB (red), and indel (green) fractions following light activation at 6 hpf in WT and MZ*polq* embryos. Full details of the kinetic model are provided in Methods. Measured data (open symbols) are overlaid with the ODE model fits (solid lines). DSB and indel are expressed as percentages of the total. The intact fraction is estimated by the model based on the DSB and indel measurements (see Fig. 3).

**Figure S5.**
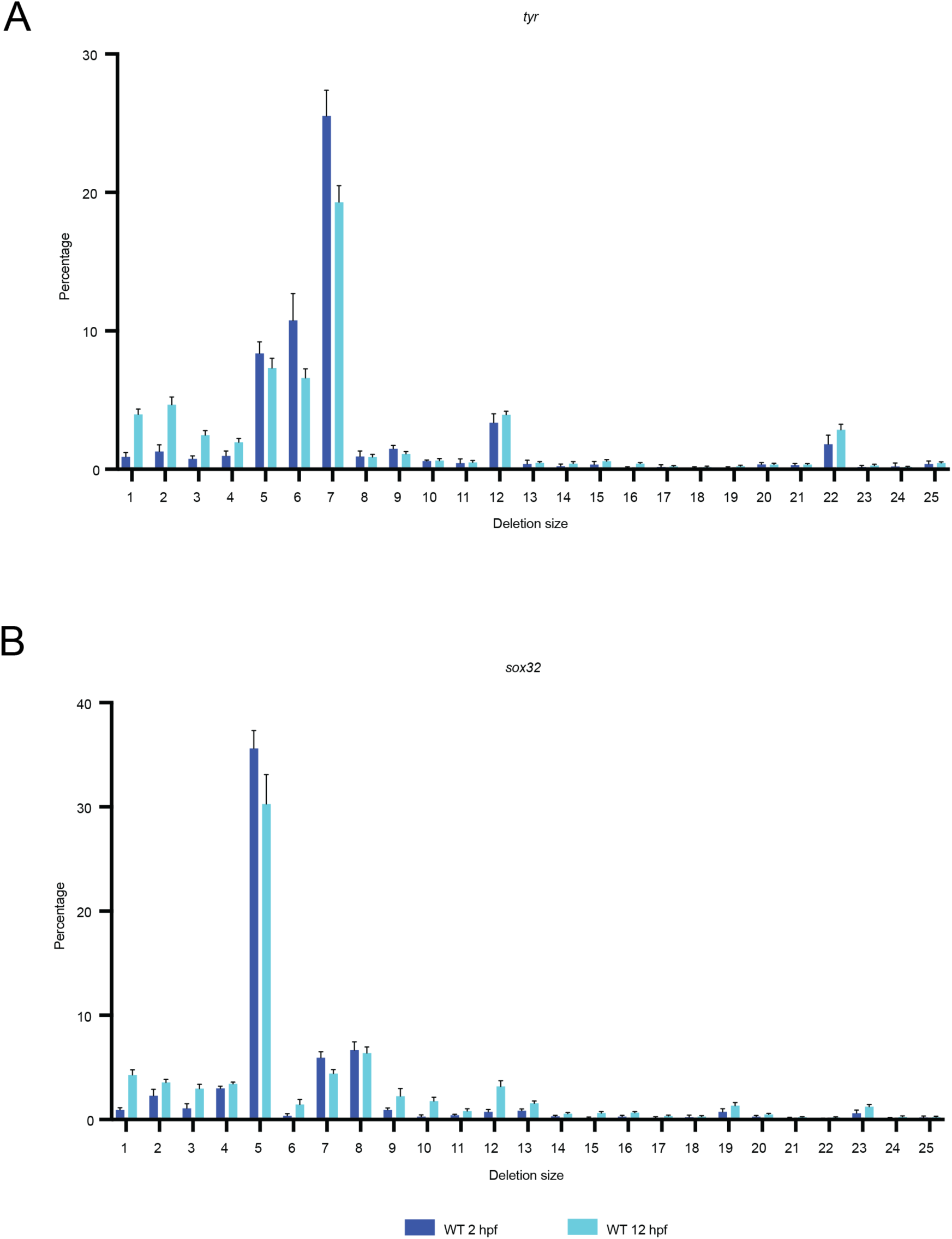
Deletion size chart at *tyr* and *sox32* loci. **A.** Mutation frequency of base pair deletions sorted by size at the *tyr* locus after light treatment in 2 and 12 hpf WT embryos. **B.** Mutation frequency of base pair deletions sorted by size at the *sox32* locus after light treatment in 2 and 12 hpf WT embryos.

**Figure S6.**
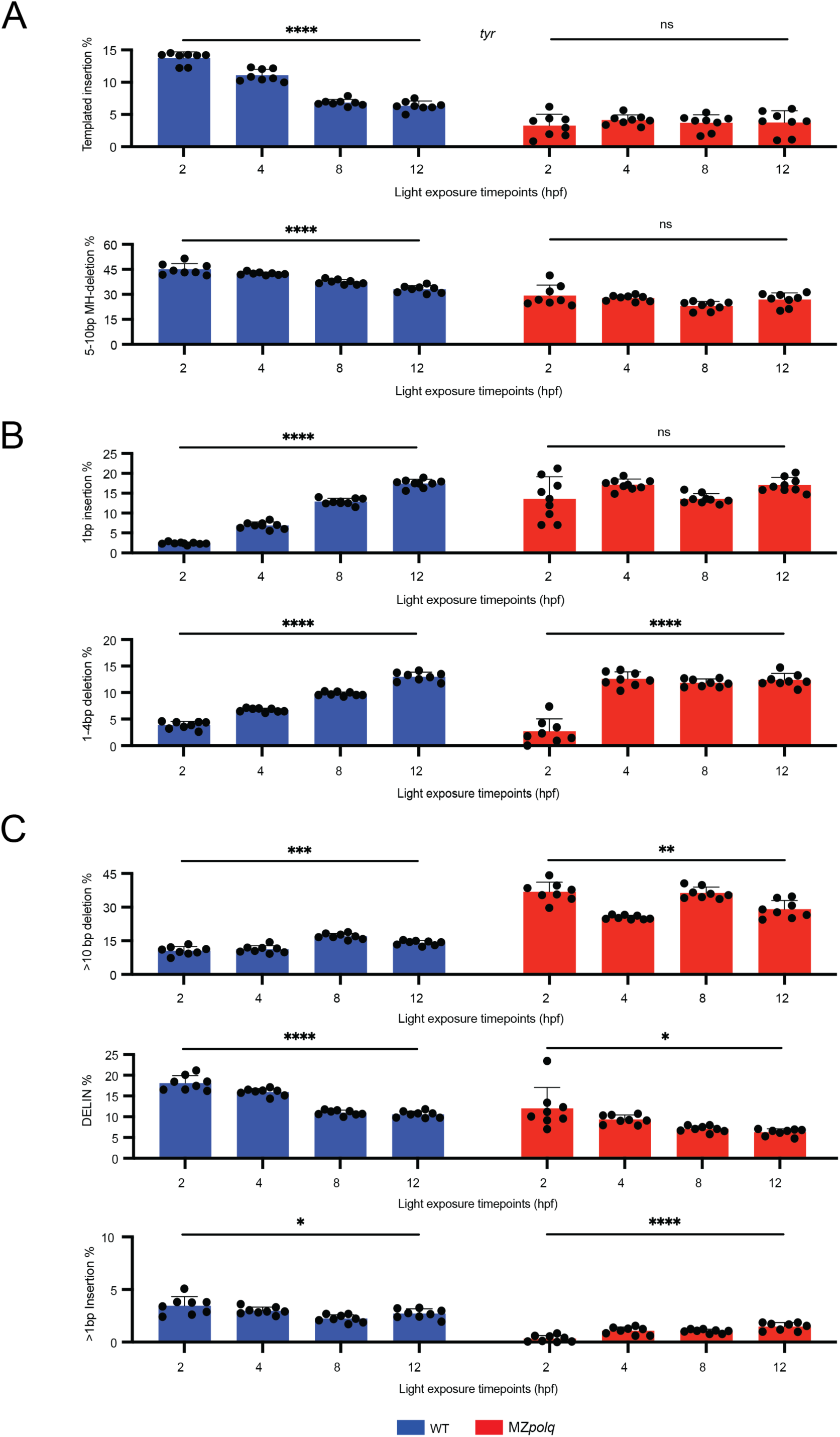
A shift in repair outcomes at the *tyr* locus as embryogenesis proceeds comparing WT and MZ*polq* embryos. **A.** Mutation frequency of MMEJ-associated repair outcomes - templated insertions (top) and microhomology-mediated 5–10 bp deletions (bottom) - at the *tyr* locus in caged *tyr* RNP-injected WT (blue) and MZ*polq* (red) embryos after light activation at the indicated time points. **B.** Mutation frequency of NHEJ-associated repair outcomes - 1 bp insertions (top) and 1–4 bp deletions (bottom, both microhomology- and non-microhomology-mediated) - at the *tyr* locus in caged *tyr* RNP-injected WT (blue) and MZ*polq* (red) embryos after light activation at the indicated time points. **C.** Mutation frequency of other repair outcomes (>10bp deletions (top), mixed deletion/insertions (DELINS)(middle), and >1bp insertions (bottom)) at the *tyr* locus in caged *tyr* RNP-injected WT (blue) and MZ*polq* (red) embryos after light activation at the indicated time points. All embryos were collected at 26–28 hpf for amplicon sequencing. Each dot represents a single embryo. Error bars in all plots represent standard deviation between 8 biological replicates. * = p ≤ 0.05, ** = p ≤ 0.01, *** = p ≤ 0.001, **** = p ≤ 0.0001, ns = not statistically significant.

**Figure S7.**
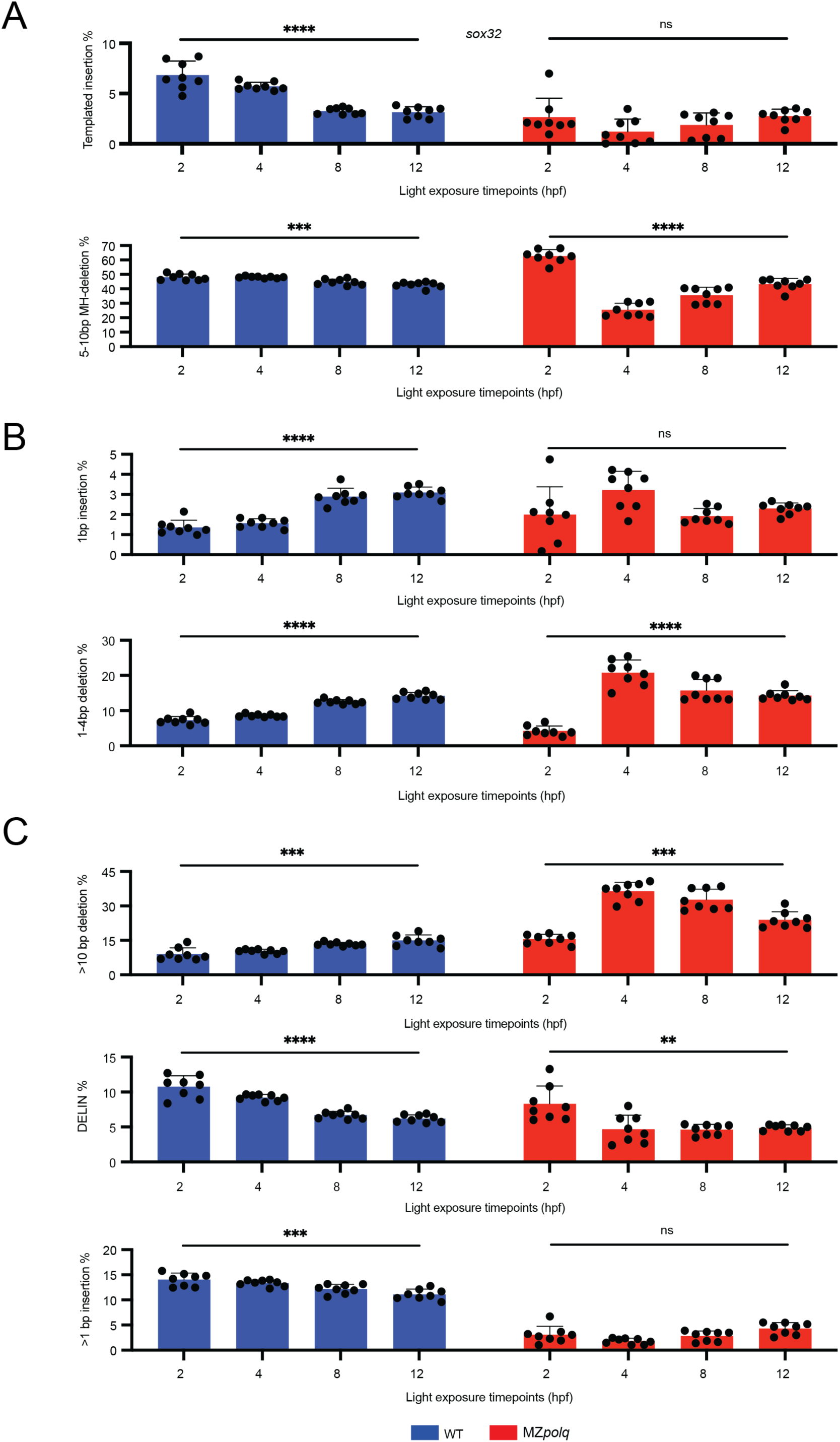
A shift in repair outcomes at the *sox32* locus as embryogenesis proceeds comparing WT and MZ*polq* embryos. **A.** Mutation frequency of MMEJ-associated repair outcomes - templated insertions (top) and microhomology-mediated 5–7 bp deletions (bottom) - at the *sox32* locus in caged *sox32* RNP-injected WT (blue) and MZ*polq* (red) embryos after light activation at the indicated time points. **B.** Mutation frequency of NHEJ-associated repair outcomes - 1 bp insertions (top) and 1–4 bp deletions (bottom, both microhomology- and non-microhomology-mediated) - at the *sox32* locus in caged *sox32* RNP-injected WT (blue) and MZ*polq* (red) embryos after light activation at the indicated time points. **C.** Mutation frequency of other repair outcomes (>10bp deletions (top), mixed deletion/insertions (DELINS) (middle), and >1bp insertions (bottom)) at the *sox32* locus in caged *sox32* RNP-injected WT (blue) and MZ*polq* (red) embryos after light activation at the indicated time points. All embryos were collected at 26–28 hpf for amplicon sequencing. Each dot represents a single embryo. Error bars in all plots represent standard deviation between 8 biological replicates. ** = p ≤ 0.01, *** = p ≤ 0.001, **** = p ≤ 0.0001, ns = not statistically significant.

**Figure S8.**
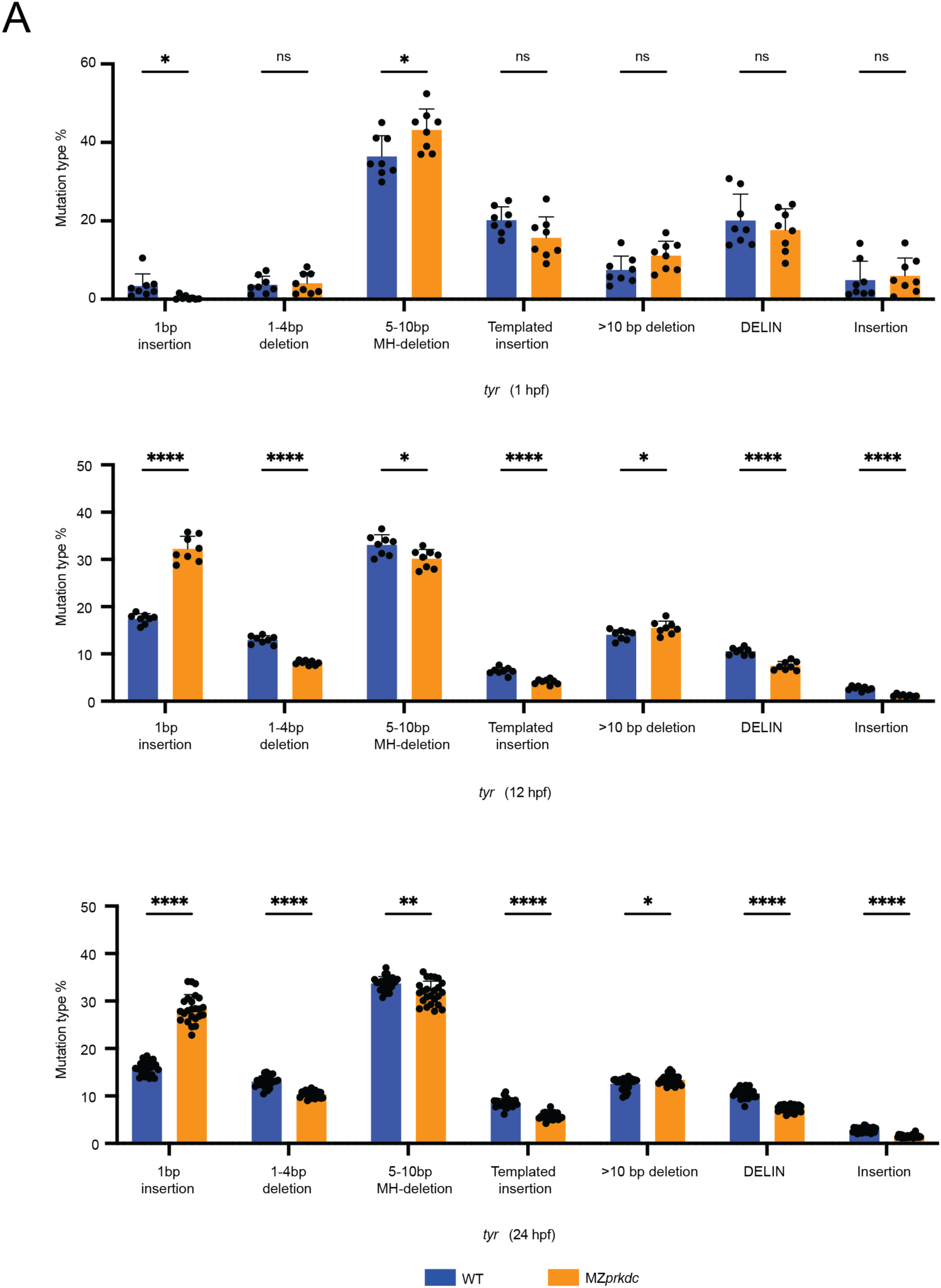
Repair outcomes at the *tyr* locus in later-stage embryos depend on NHEJ. **A.** Mutation frequency of seven repair outcomes at the *tyr* locus in caged *tyr* RNP-injected WT (blue) and MZ*prkdc* (orange) embryos after light activation at 1 hpf (top), 12 hpf (middle), or 24 hpf (bottom). Embryos after light activation at 1 and 12 hpf were collected at 26–28 hpf and embryos after light activation at 24 hpf were collected at 48 hpf for amplicon sequencing. Each dot represents a single embryo. Error bars represent standard deviation between 8 biological replicates for 1 and 12 hpf embryos and 24 biological replicates for 24 hpf embryos. * = p ≤ 0.05, ** = p ≤ 0.01, **** = p ≤ 0.0001, ns = not statistically significant.

**Figure S9.**
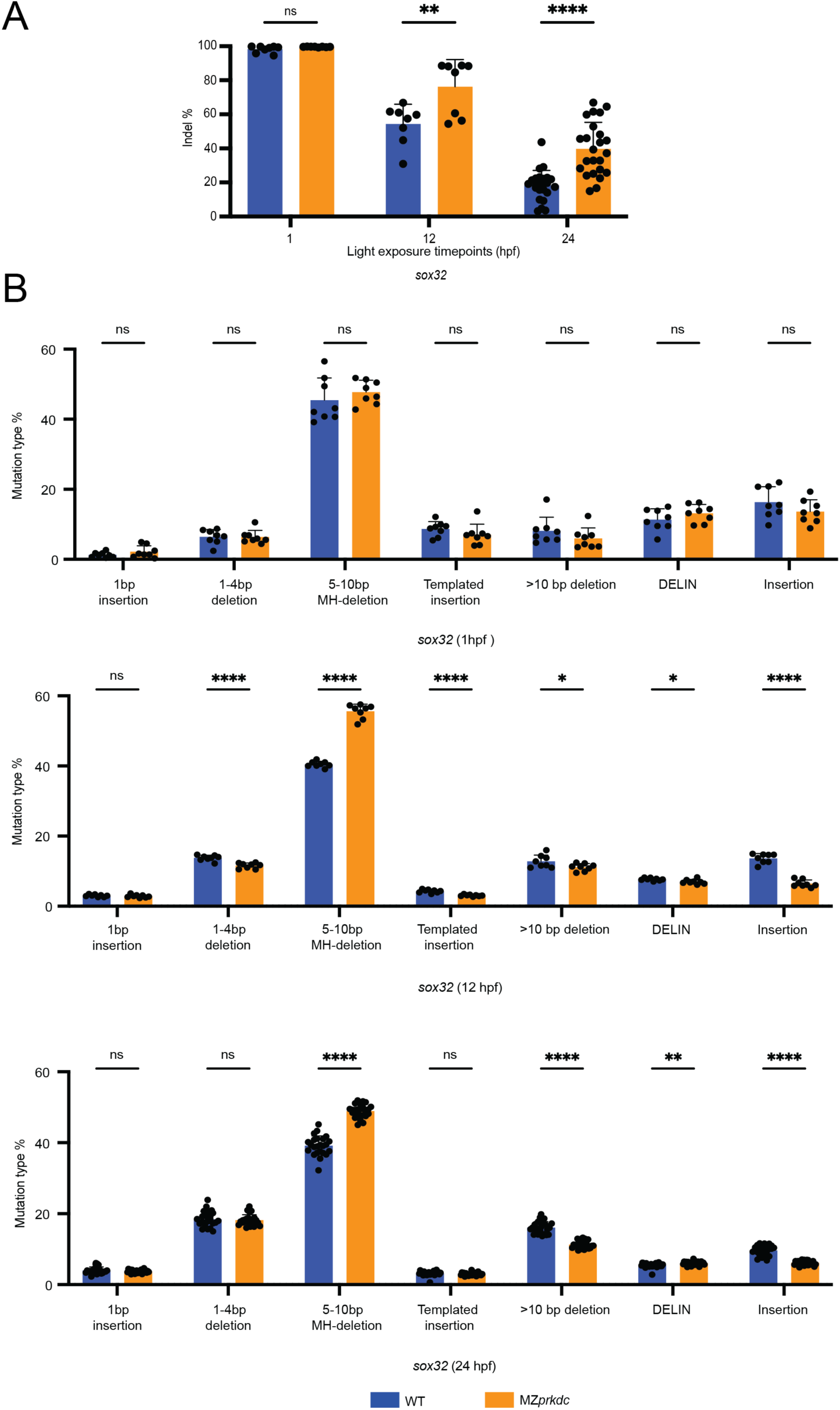
DNA repair fidelity and outcomes at the *sox32* locus in later-stage embryos depend on NHEJ. **A.** Indel frequency at the *sox32* locus in caged *sox32* RNP-injected WT (blue) and MZ*prkdc* (orange) embryos after light activation at 1, 12, or 24 hpf. **B.** Mutation frequency of seven repair outcomes at the *sox32* locus in caged *sox32* RNP-injected WT (blue) and MZ*prkdc* (orange) embryos after light activation at 1 hpf (top), 12 hpf (middle), or 24 hpf (bottom). Embryos after light activation at 1 and 12 hpf were collected at 26–28 hpf and embryos after light activation at 24 hpf were collected at 48 hpf for amplicon sequencing. Each dot represents a single embryo. Error bars represent standard deviation between 8 biological replicates for 1 and 12 hpf embryos and 24 biological replicates for 24 hpf embryos. * = p ≤ 0.05, ** = p ≤ 0.01, **** = p ≤ 0.0001, ns = not statistically significant.

**Table S1.**
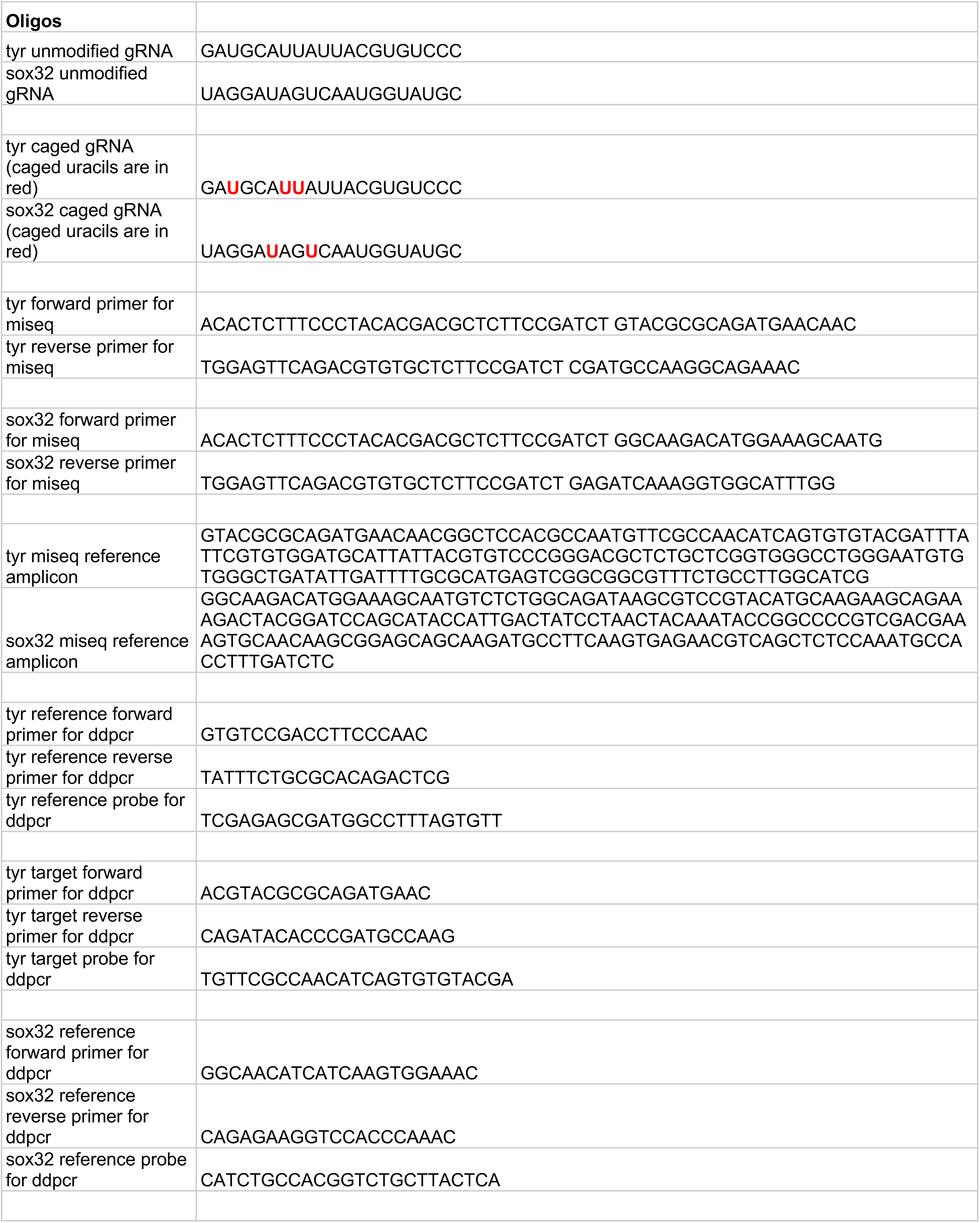
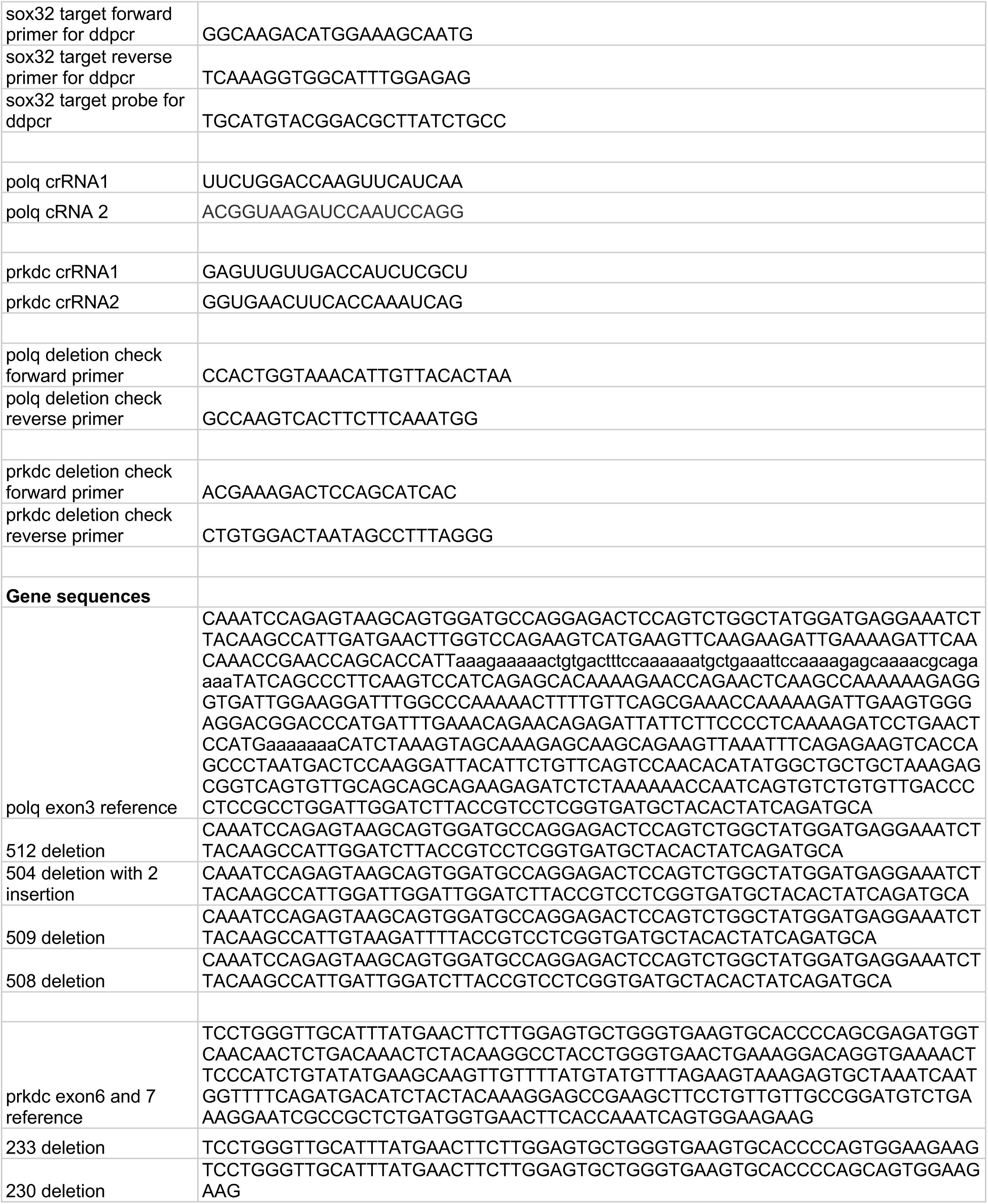
DNA sequences.

**Table S2.** Parameters from kinetics model.

|  |  |  |  |  |
| --- | --- | --- | --- | --- |
| Fitting parameters obtained from kinetics model |  |  |  |  |
| Parameter | WT 3 hpf | MZpolq 3 hpf | WT 6 hpf | MZpolq 6 hpf |
| kc | 0.0231±0.0051 | 0.0231±0.0051 | 0.0187±0.0037 | 0.0187±0.0037 |
| kp | 0.0064±0.0060 | 0.0057±0.0037 | 0.0098±0.0041 | 0.0091±0.0034 |
| km | 0.0135±0.0021 | 0.0026±0.0008 | 0.0038±0.0008 | 0.0010±0.0006 |
| B0 | 0.2086±0.0423 | 0.1637±0.0439 | 0.1709±0.0345 | 0.1605±0.0349 |
| R2 | 0.9634 | 0.9634 | 0.9773 | 0.9773 |

